# Integrated single-cell chromatin and transcriptomic analyses of human scalp reveal etiological insights into genetic risk for hair and skin disease

**DOI:** 10.1101/2022.09.28.509620

**Authors:** Benjamin Ober-Reynolds, Chen Wang, Justin M. Ko, Eon J. Rios, Sumaira Z. Aasi, Mark M. Davis, Anthony E. Oro, William J. Greenleaf

## Abstract

Genome-wide association studies (GWAS) of hair and skin disease have identified many genetic variants associated with disease phenotypes, but identifying causal variants and interpreting their function requires deciphering gene-regulatory networks in disease-relevant cell types. To this end, we generated matched scRNA- and scATAC-seq profiles of human scalp biopsies, identifying diverse cell types of the hair follicle niche. By interrogating the integrated datasets across multiple levels of cellular resolution, we infer 50-100% more enhancer-gene links than prior approaches, and show that the aggregate accessibility at linked enhancers for highly-regulated genes predicts expression. We use these gene-regulatory maps to prioritize cell types, genes, and causal variants implicated in the pathobiology of androgenetic alopecia (AGA), eczema, and other complex traits. AGA GWAS signals are strongly and specifically enriched in dermal papilla cell open chromatin regions, supporting the role of these cells as key drivers of AGA pathogenesis. Finally, we trained machine-learning models to nominate SNPs that affect gene expression through disruption of specific transcription factor binding motifs, identifying candidate functional SNPs linked to expression of WNT10A in AGA and IL18RAP in eczema. Together, this work reveals principles of gene regulation and identifies gene regulatory consequences of natural genetic variation in complex skin and hair diseases.

## Introduction

Skin is composed of a complex, interconnected community of cell types from diverse developmental origins that perform coordinated functions to enable tissue homeostasis. For example, skin contains hair follicles– dynamic mini-organs that progress through well-defined cycles of growth (anagen), regression (catagen), and resting (telogen), guided by paracrine signals from their surrounding stromal and immune niche to the stem cell reservoir^1–3^. Disruption of these interconnected cellular communities cause human skin and hair diseases such as alopecia areata (AA), in which normal hair follicle cycling is prevented by auto-reactive T-cells, or androgenetic alopecia (AGA), in which hair follicles gradually miniaturize as a result of a poorly understood interplay of genetic and hormonal factors. Understanding the pathobiology of these and other skin and hair diseases thus requires approaches capable of identifying perturbations across a multitude of potentially relevant cell types and states.

While genome-wide association studies (GWAS) have identified dozens of distinct genomic loci associated with complex hair and skin-related disorders^4–8^, identifying specific causal variants and interpreting their molecular function is rarely straightforward. Indeed, the majority of GWAS disease risk variants reside in non-coding genomic regions, and many are predicted to exert their effects through disruption of the regulatory function of these cis-regulatory elements^9^. These cis-regulatory elements tend to be highly cell-type specific, and in many cases do not exert their effects on the nearest gene. Identifying causal variants and interpreting their function thus requires deciphering gene-regulatory networks in disease-relevant cell types.

Single-cell genomics, primarily single-cell RNA (scRNA) sequencing, has begun to allow for the identification and characterization of the diverse cell types in human skin in healthy and disease contexts ^10–16^. However, to date these studies have focused primarily on isolation of specific cell lineages, or use sample collection strategies where recovery of complete accessory structures, such as hair follicles, is not achieved. Furthermore, while scRNA sequencing studies provide a snapshot of the transcriptional state of cell types within a tissue, the underlying cis-regulatory DNA elements that program these expression states are not observed, precluding deeper insights into the distal elements that drive these gene expression states and how non-coding genetic variation at these elements influences disease phenotypes.

In this study, we expand the current understanding of gene regulatory networks in healthy and diseased human skin and hair follicles by generating paired, single-cell atlases of gene expression and chromatin accessibility in primary human scalp from healthy controls and from patients with alopecia areata. We use these integrated multi-omic landscapes to identify enhancer-gene linkages at multiple scales of cellular resolution, yielding 50-100% more enhancer-gene links than similarly powered prior multi-omic studies. We identify a subset of key cell lineage genes associated with a disproportionately large number of cis-regulatory elements, and show that the expression of many of these highly regulated genes (HRGs) is driven by distinct combinations of enhancer modules across a range of cell types, with increasing enhancer recruitment resulting in higher transcript levels. We use our integrated multi-omic data to define the chromatin and gene expression landscapes of interfollicular and hair follicle-associated keratinocyte subpopulations, and to predict regulatory gene targets of transcription factors (TFs) driving distinct keratinocyte differentiation trajectories. Finally, we perform integrative analyses using our atlas of scalp regulatory elements and existing GWAS data sets to identify relevant cell types and putative target genes for fine-mapped SNPs associated with skin and hair diseases. GWAS signals associated with AGA were strongly and specifically enriched in dermal papilla open chromatin regions, and linked to target genes enriched for roles in WNT-signaling. We further extend these analyses by training machine-learning models to nominate identifying 47, 19, and 19 prioritized SNPs for AGA, eczema, and hair color respectively based on their predicted effects on cell-type specific chromatin accessibility.

## Results

### A paired transcriptomic and epigenetic atlas reveals cellular diversity of human scalp

To define the cellular heterogeneity and gene-regulatory complexity of human scalp, we created paired, single-cell transcriptomic and chromatin accessibility atlases from healthy and disease-affected primary human scalp tissue. We obtained full-thickness human scalp tissue from three sources: punch biopsy samples from healthy control volunteers (C_PB, n = 3), punch biopsies from patients with clinically active alopecia areata (AA, n = 5), and discarded normal peripheral surgical tissue from patients undergoing dermatologic surgeries on the scalp (C_SD, n = 7) (Table S1, Figure 1A,B). For each of these samples, we dissociated scalp tissue into single cell suspensions and prepared single-cell RNA sequencing (scRNA-seq) and single-cell ATAC sequencing (scATAC-seq) libraries using the Chromium platform from 10x Genomics. In total, we obtained 54,288 single-cell transcriptomes and 45,896 single-cell chromatin accessibility profiles after stringent quality control and filtering of each dataset (Methods, Figure S1A-C). Using an Iterative LSI dimensionality reduction and clustering approach ^17^, we identified 22 cell clusters from scRNA-seq data (Figure 1C). Using a similar dimensionality reduction and clustering approach for scATAC-seq data, we identified 22 distinct cell clusters (Figure 1D).

**Figure 1.**
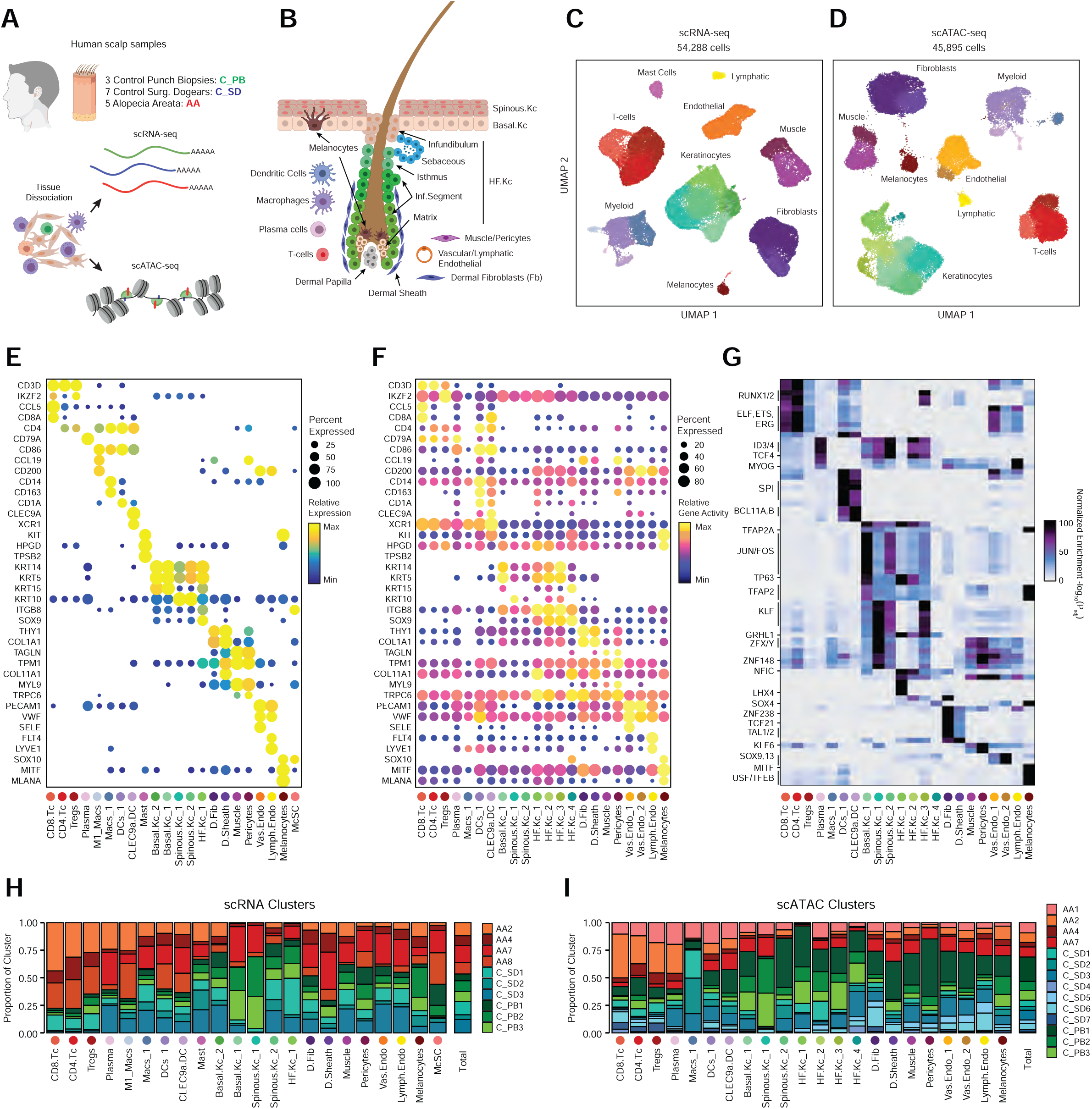
Multi-omic single-cell atlas of primary human scalp (A) Samples and profiling methods used in this study. (B) Schematic representation of the cellular diversity within human scalp. (C, D) UMAP representation of all scRNA-seq (C) and scATAC-seq (D) cells passing quality control, colored by annotated clusters. Broad cell types are labeled on the UMAP and higher resolution labels are shown in (E-F). (E) scRNA gene expression for selected marker genes for each scRNA-seq cluster. The color indicates the relative expression across all clusters and the size of the dot indicates the percentage of cells in that cluster that express the gene. (F) scATAC gene activity scores for the same markers shown in (E). (G) Hypergeometric enrichment of transcription factor motifs in marker peaks for each scATAC-seq cluster. (H) The fraction of each sample comprising each scRNA-seq cluster. Samples from control punch biopsies are shown in shades of green, surgical dogears in shades of blue, and patients with alopecia areata shown in red. The total proportions for each sample are shown in the rightmost column. (I) Same as (H), but for the scATAC-seq clusters.

To annotate clusters in the scRNA-seq, we examined the gene expression of known marker genes (Figure 1E, S1F). To annotate clusters in the scATAC-seq data, we used ‘gene activity scores’ – a gene-level metric aggregating the chromatin accessibility around a gene-body – for known marker genes (Figure 1F, S1G)^17, 18^. For example, interfollicular basal keratinocyte clusters exhibited high gene activity and expression of the basal keratin KRT15 ^19^, hair-follicle (HF) keratinocyte clusters exhibited high gene activity and expression of the transcription factor SOX9 ^20^, T-lymphocyte clusters exhibited high gene activity and expression of the cell surface marker CD3D, and fibroblast clusters exhibited high gene activity and expression of the cell surface marker THY1 ^21^. We observed a relatively large scRNA-seq cluster expressing high levels of mast-cell markers including beta tryptases (TPSB1/2) and HPGD ^22–24^, but we did not observe a corresponding scATAC-seq cluster, perhaps due to the tendency for granulocyte chromatin to spontaneously decondense during nuclear isolation ^25, 26^. To better resolve the cellular heterogeneity of broad cell groupings, we sub-clustered 5 major cell classes (keratinocytes, T-lymphocytes, myeloid lineage cells, fibroblasts, and endothelial cells) in both the scRNA- and scATAC-seq datasets (Methods, Figure S2A-C). We again used the gene expression and gene activity scores of known marker genes to label clusters in the sub-clustered scRNA- and scATAC-seq data respectively (Figure S2C). This higher-resolution level of clustering revealed several rare cellular subtypes that were not well resolved at the full dataset level, such as dermal papilla cells (HHIP, WNT5A, PTCH1) in the fibroblast cell group ^27^, eccrine gland cells (AQP5, KRT19) in the keratinocyte cell group ^28^, and TREM2 positive macrophages (TREM2, OSM) in the myeloid cell group, which have been described as having a role in hair follicle cycle regulation in mice ^29^. This high-resolution clustering approach identified a total of 42 scRNA-seq (10 keratinocyte, 7 fibroblast, 6 endothelial, 6 T-lymphocyte, 8 myeloid, and 5 other) and 38 scATAC-seq (11 keratinocyte, 6 fibroblast, 5 endothelial, 5 T-lymphocyte, 7 myeloid, and 4 other) clusters, hereafter referred to as ‘high-resolution clusters.’ We observed excellent overall correspondence between high-resolution cluster cell types and marker genes between the two modalities (Figure 1, S1, S2). We used the high-resolution scATAC-seq clusters to identify a union set of 589,294 ‘peaks’ of open chromatin corresponding to cis-regulatory elements across the entire dataset (Methods) ^18^. We identified 182,498 peaks that were differentially accessible between the broad scATAC clusters (Wilcoxon FDR ≤ 0.1; Log2FC ≥ 0.5) (Figure S1H), and found that these cluster-specific peaks were enriched for motifs of transcription factors that drive cell lineages, such as RUNX and ETS factors in T-lymphocytes ^30, 31^, SPI (PU.1) factors in myeloid lineage cells ^32^, TP63 in keratinocytes ^33^, and MITF in melanocytes ^34^ (Figure 1G).

For both the scRNA- and scATAC-seq datasets, all clusters were composed of cells spanning the majority of patient donors, and cells originating from distinct patient donors were well-mixed within each cluster (Figure S1D,E). Certain cell types were more abundant in certain samples groups however: T-lymphocyte clusters comprised a larger fraction of cells obtained from alopecia areata patient samples (AA) and certain HF keratinocyte clusters were depleted in these same samples (Figure 1H,I, S1I,J). These observations align with current understanding of alopecia areata pathophysiology, which involves peribulbar HF T-cell infiltration and disruption of normal HF cycling ^35^.

#### Highly regulated genes use distinct enhancer modules to tune gene expression

After annotation of low- and high-resolution clusters in both the scRNA- and scATAC-seq datasets, we bioinformatically integrated our scATAC and scRNA datasets using canonical correlation analysis (CCA) ^36^. We computed Jaccard indices between the scATAC and scRNA cluster annotations and observed high correspondence between cell types in the full dataset and in each of the sub-clustered datasets (Figure S2D,E). We used these integrated scATAC and scRNA datasets to identify cis-regulatory elements with accessibility correlated to local gene expression (‘peak-to-gene links’) by performing a bias-corrected correlation analysis between expressed genes and peaks in their broad genomic vicinity (Methods) ^17, 18^. To enhance our ability to detect peak-to-gene linkages relevant to both broad cell type identity and to regulation of more similar cell subtypes, we performed dataset integration and peak-to-gene linkage identification on the full scalp dataset, as well as on each of the sub-clustered datasets (keratinocytes, T-lymphocytes, myeloid lineage cells, fibroblasts, and endothelial cells) (Figure 2A). Across all datasets, we identified 146,088 unique peak-to-gene links (Correlation ≥ 0.5, Figure S2F), observing that genes had a median of four linked peaks (Figure S2G). Only 66,702 of these links were detected using the full, non-sub clustered dataset (Figure S2F), but the 79,386 additional peak-to-gene linkages identified from this hierarchical linkage approach were as well supported as the links identified on the full dataset– peak-to-gene linkages detected in the full dataset and in each of the sub-clustered datasets were all more likely to be evolutionarily conserved than unlinked peaks (Figure S2H, Wilcoxon rank-sum test p < 2.2 x 10^-16^ for each comparison), and links identified on each of the sub-clustered datasets were significantly more likely to be corroborated by cell-type specific enhancer-gene pair predictions in a recent 131 human cell type and tissue activity-by-contact (ABC) model dataset when compared to distance-matched, permuted peak-to-gene linkages (Figure S2I, Hypergeometric enrichment test p < 2.2 x 10^-16^ for each comparison)^37, 38^. The majority of peaks (491,106; 83.3%) were not linked to any gene in this analysis, consistent with the relatively small expected effect size of most cis-regulatory elements ^39^. To demonstrate how enhancer-gene links can be missed by using only the full dataset, we examined peaks linked to RUNX3, a gene that is expressed in multiple distinct cell types. Using the full, non-sub clustered dataset, we identified 43 linked peaks for RUNX3, the majority of which were accessible in T-lymphocytes, myeloid lineage cells, and melanocytes. Relatively little chromatin accessibility was observed in keratinocytes (Figure 2B). However, performing peak-to-gene identification in the sub-clustered keratinocytes revealed numerous RUNX3-linked peaks that were strongly accessible in sebaceous gland cells. Repeating this process on each of the sub-clustered data sets resulted in a non-redundant set of 81 RUNX3-linked peaks, nearly doubling the number of RUNX3-linked enhancers identified using only the full dataset.

**Figure 2.**
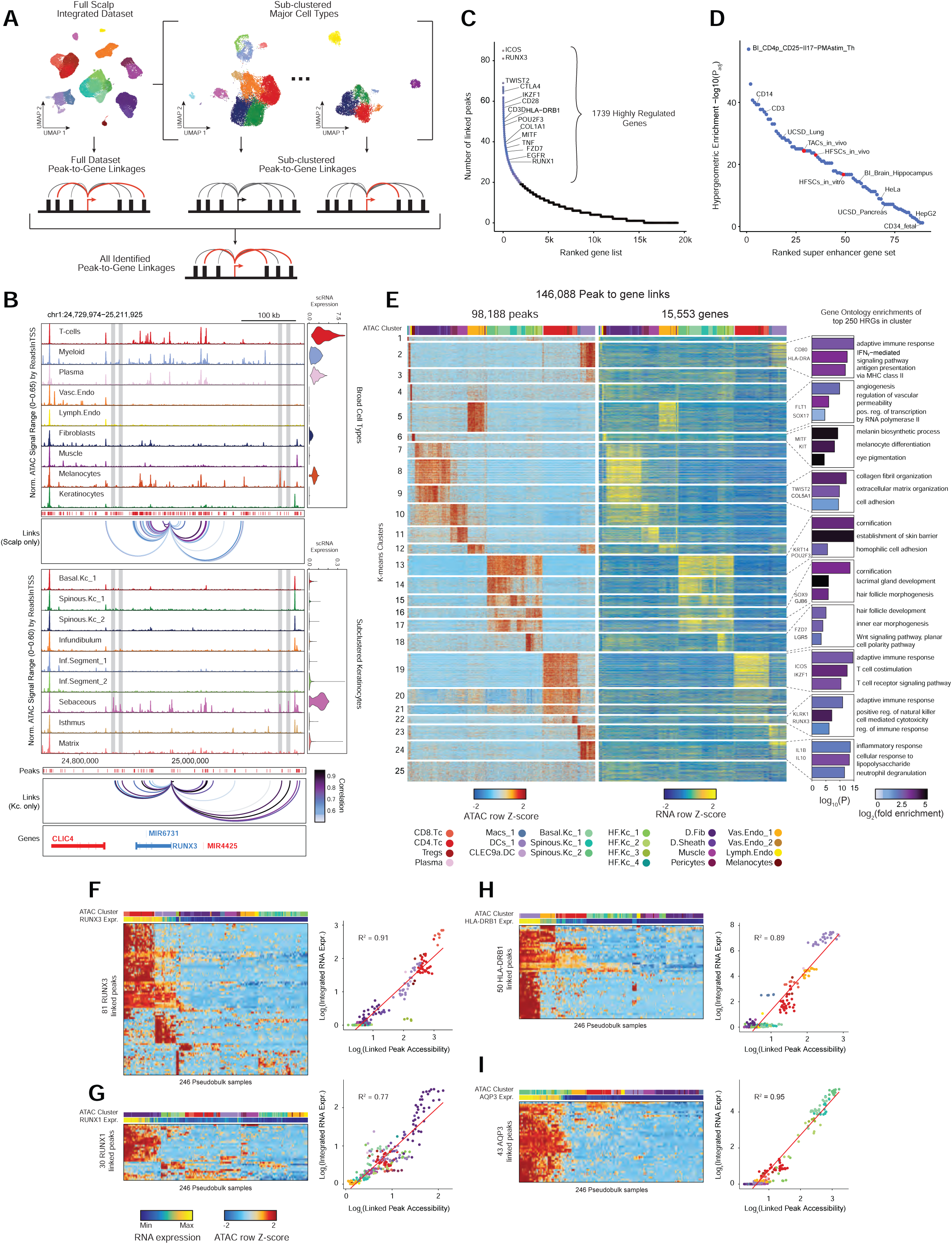
Gene regulatory dynamics and modularity in human scalp (A) Peak-to-gene linkages were identified on the integrated scATAC and scRNA full scalp dataset, and on each of the five major cell-type sub-clustered datasets. Peak-to-gene linkages identified in each dataset are merged to form the full set of peak-to-gene linkages. (B) Genomic tracks for accessibility around RUNX3 for different cell types. Integrated RUNX3 expression levels are shown in the violin plot for each cell type to the right. Loops shown below the top panel indicate peak-to-gene linkages identified on only the full integrated dataset. Lower panel shows the genomic tracks for accessibility around RUNX3 for sub-clustered keratinocytes. Loops shown below these tracks indicate peak-to-gene linkages identified on the sub-clustered dataset. (C) Genes ranked by the number of peak-to-gene links identified for each gene. 1739 ‘highly regulated genes’ (HRGs) had ≥ 20 peak-to-gene linkages. (D) Hypergeometric enrichment of super-enhancer linked genes in human scalp HRGs for each cell and tissue type profiled in Hnisz et al. 2013. Red dots represent enrichment of the human homolog of mouse hair-follicle super-enhancer linked genes identified in Adam et al. 2015. (E) Heatmap showing the chromatin accessibility (left) and gene expression (right) for the 146,088 peak-to-gene linkages. Peak-to-gene linkages were clustered using k-means clustering (k = 25). Sample top HRGs for select clusters are shown to the right of the gene expression heatmap. GO term enrichments for the top 200 genes ranked by number of peak-to-gene linkages are shown to the right for select k-means clusters. (F) Heatmap showing chromatin accessibility at RUNX3-linked peaks for 246 pseudo-bulked scATAC-seq samples. Cell type labels are shown in the bar above the heatmap, and RUNX3 expression levels for each pseudo-bulk are shown below. Scatter plot to the right shows the relationship between linked peak accessibility and resulting gene expression for each of the pseudo-bulked samples shown in the heatmap to the left. (G) Same as in (F), but for RUNX1. (H) Same as in (F), but for HLA-DRB1. (I) Same as in (F), but for AQP3.

Consistent with other paired scRNA- and scATAC-seq studies, we identified a subset of genes that were associated with notably greater numbers of putative enhancers than the majority of expressed genes ^40, 41^. We identified 1,739 such “highly regulated genes” (HRGs) across all scalp cell types by ranking genes by their number of linked peaks and retaining those that exceeded the inflection point at 20 peak-to-gene linkages (Figure 2C). These genes include transcription factors driving cell identity, such as RUNX1, TWIST2, and MITF, as well as genes associated with critical cell type-specific functions, such as COL1A1, KRT14, and ICOS. HRGs identified in scalp were also significantly enriched for previously identified “super-enhancer” associated genes from a range of tissues and cell lines (Figure 2D) ^42, 43^. To explore the regulatory heterogeneity of HRGs across different cell types, we clustered pseudo-bulked samples by the accessibility of gene-linked cis-regulatory elements, using k-means clustering to identify co-occurring regulatory modules (Figure 2E). Within these K-means clusters, we again ranked genes by the number of linked enhancers and performed gene ontology (GO)-term enrichment on the top 250 genes in each cluster. HRGs from each cluster were strongly enriched for cell-type specific GO-terms, including ‘adaptive immune response’ (Myeloid clusters), ‘melanocyte differentiation’ (melanocytes), and ‘hair follicle development’ (HF keratinocytes) (Figure 2E).

While many HRGs are specifically expressed in one or a few closely related cell types, several HRGs, such as RUNX3, were expressed in multiple distinct cell types and associated with distinct chromatin peaks in different cellular contexts (Figure 2B). To explore the cis-regulatory element heterogeneity of single HRGs across multiple cell types, we hierarchically clustered pseudo-bulk samples according to the accessibility observed in all peaks linked to a single HRG (Methods). For many HRGs, we observed multiple ‘modules’ of co-accessible cis-regulatory elements active in distinct cell types (Figure 2F-I). These modules contained variable numbers of cis-regulatory elements, and some modules were shared by multiple cell types while others were highly specific to a single cell type. Interestingly, the aggregate accessibility signal observed across all linked peaks was strongly correlated with the gene expression of the linked gene (Figure S2J), suggesting that aggregate chromatin accessibility at linked cis-regulatory elements provides a proxy for gene expression, irrespective of which specific cis-regulatory elements are active. These findings support a broadly additive, modular model of enhancer activity– a model that has been supported by genetic perturbation studies of individual enhancer elements of alpha-globin^44^ and Myc^45^, studies of enhancers involved in limb development^46^, and through genomic-scale measures of enhancer activity ^47, 48^.

#### Gene-regulatory diversity of interfollicular and follicular keratinocytes

While the transcriptional heterogeneity of human interfollicular^11, 16^ and follicular^49^ keratinocytes are well known, our peak-to-gene linkage analysis enables a deeper interrogation of the cell type regulatory logic of primary human skin and hair. Our high-resolution sub-clustered keratinocyte dataset comprised 10 total high-resolution clusters, with three clusters of interfollicular keratinocytes (Basal.Kc_1, Spinous.Kc_1 and _2), six hair follicle associated clusters (Infundibulum, Isthmus, Sebaceous, Inf.Segment_1 and _2, Matrix), and one small cluster corresponding to eccrine gland cells (Eccrine) (Figure 3A-B, S3A-B). To focus on the differences in gene-regulatory mechanisms between keratinocyte subsets, we used peak-to-gene linkages specific to keratinocyte cell types (28,991 links, correlation > 0.45) for subsequent analyses. K-means clustering of these linkages revealed extensive gene-regulatory diversity within keratinocyte subtypes, with co-accessible peaks in these clusters strongly enriched for distinct transcription factor motifs (Figure S3C,D). To identify TFs that play a significant regulatory role in specifying distinct keratinocyte subsets, we first identified motifs with variable accessibility between keratinocyte populations (Figure S3E, y-axis). To differentiate between TFs from the same family that share similar motifs, we additionally correlated each TF’s gene expression with its motif activity across the diversity of keratinocyte cell types (Figure S3E, x-axis). This approach identified TFs previously described as having important roles in skin (TP63, FOSL1, KLF4)^33, 50, 51^ and hair differentiation (SOX9, LHX2, HOXC13)^43, 52, 53^. Some TFs were broadly active in multiple related cell types, such as TP63 in basal keratinocyte clusters, and SOX9 in HF keratinocyte clusters, while others were more cell type-specific– LHX2 motifs were highly accessible only in the cycling, inferior segment of the hair follicle, and RUNX3 motifs were specifically accessible in a distinct population of sebaceous gland cells (Figure 3C).

**Figure 3.**
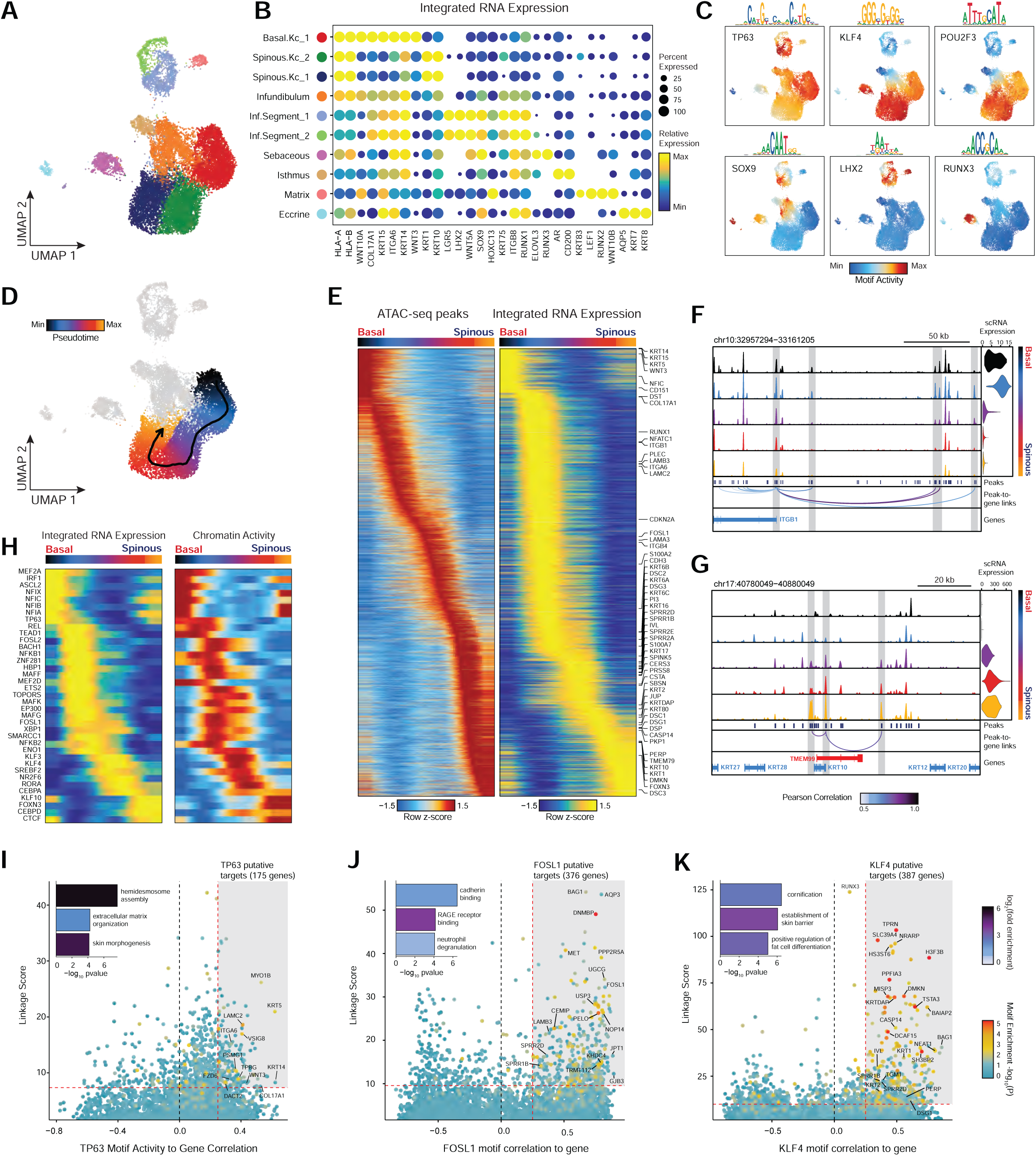
Scalp keratinocyte diversity and regulatory control of interfollicular keratinocyte differentiation. (A) UMAP representation of subclustered keratinocytes in the scATAC-seq dataset (B) Integrated gene expression for select markers across keratinocyte subtypes. The color indicates the relative expression across all clusters and the size of the dot indicates the percentage of cells in that cluster that express the gene. (C) ChromVAR deviation z-scores showing transcription factor motif activity for select transcription factors. (D) Slingshot differentiation trajectory starting with basal IFE keratinocytes and progressing to upper layer spinous keratinocytes. (E) Heatmap of 10% most variable peaks (n = 31,333) and 10% most variable genes (n = 2,127) along the trajectory from basal to spinous keratinocytes. (F) Genomic tracks of accessibility around the ITGB1 promoter. Tracks are pseudo-bulked samples ordered along the IFE differentiation trajectory. Integrated ITGB1 expression levels are shown in the violin plot for each pseudo-bulk to the right. (G) Same as in (F), but for the KRT10 promoter. (H) Paired heatmaps of positive TF regulators whose TF motif activity (left) and matched gene expression (right) are positively correlated across the interfollicular keratinocyte differentiation pseudotime trajectory. (I) Prioritization of gene targets for TP63. The x-axis shows the Pearson correlation between the TF motif activity and integrated gene expression for all expressed genes across all keratinocytes. The y-axis shows the TF Linkage Score (for all linked peaks, sum of motif score scaled by peak-to-gene link correlation). Color of points indicates the hypergeometric enrichment of the TF motif in all linked peaks for each gene. Top gene targets are indicated in the shaded area (motif correlation to gene expression >0.25, linkage score >80th percentile). GO term enrichments for the top gene targets are shown in the inset bar plot. (J) Same as in (I), but for FOSL1. (K) Same as in (I), but for KLF4.

#### Identification of regulatory gene targets of transcription factors driving interfollicular keratinocyte differentiation

Interfollicular keratinocytes undergo continuous replacement by coordinated differentiation and outward migration of basal keratinocytes to spinous, granular, and finally cornified keratinocytes. To identify TF drivers of this differentiation pathway in a human *in vivo* context, we constructed a semi-supervised pseudotemporal trajectory between basal interfollicular keratinocytes and more differentiated spinous keratinocytes (Methods) (Figure 3D). Visualization of the most variable 10% of peaks along this trajectory revealed a continuous, gradual opening and closing of accessible chromatin (Figure 3E). The most variable 10% of genes along this pseudotemporal ordering included known transcriptional changes that occur during interfollicular keratinocyte terminal differentiation, with cells earliest in the trajectory expressing basal keratins (KRT15, KRT5, KRT14) and hemidesmosome components (ITGA6, ITGB1, COL17A1), and cells later in the trajectory expressing suprabasal keratins (KRT1 and KRT10) (Figure 3E) ^54^. Genomic tracks of the ITGB1 gene locus, active in basal keratinocytes, and KRT10, active in spinous layer keratinocytes, demonstrate how changes in accessibility at linked enhancers for each gene correlates with changes in gene expression across differentiation (Figure 3F,G). To identify TF drivers of interfollicular keratinocyte differentiation, we correlated TF motif activity with expression using only cells along the interfollicular keratinocyte differentiation trajectory (Figure 3H). This analysis identified several TFs with previously identified sequential roles in interfollicular keratinocyte differentiation, such as TP63 activity in early-mid differentiation followed by KLF3/4, RORA, and CEBPA/D in later differentiation^50, 55^.

We next sought to identify potential regulatory gene targets of TFs involved in driving keratinocyte cell identity. For TFs previously identified as potential regulatory drivers (Figure 3H, S3E) we correlated the TF’s motif activity with the integrated gene expression of all expressed genes across pseudo-bulked samples. Next, for each gene, we selected all linked peaks that contained an instance of the TF motif and computed a ‘linkage score’ that aggregates the strength of the peak-to-gene linkage and the confidence of its motif match for all linked peaks (Methods). Finally, we computed the hypergeometric enrichment of TF motif matches in peaks linked to that gene. Using this approach, we identify potential regulatory targets of a TF as genes that are both highly correlated with global motif activity and have a high linkage score specifically for that TF (Pearson correlation > 0.25 and linkage score > 80th percentile). We identified 175 genes as potential direct TP63 regulatory targets (Figure 3I). We found that these predicted TP63 regulatory targets were enriched for genes that were downregulated (OR = 1.95, Fisher’s exact test p-value = 0.0002), but not upregulated (OR = 0.71) in keratinocytes with inactivating TP63 mutations ^56^. These regulatory targets included basal keratins KRT5 and KRT14, as well as genes involved in the anchoring of keratinocytes to the basement membrane, such as LAMC2, ITGA6, and COL17A1. When we performed GO-term enrichment analysis on these 175 genes, the most significant annotation was for genes associated with hemidesmosome assembly, consistent with TP63’s known role in the regulation of cell adhesion ^57^. FOSL1, a factor active in the intermediate stages of differentiation, instead was linked to gene targets enriched for cadherin binding functionality, a key regulatory signal in early keratinocyte differentiation (Figure 3J)^58^. For KLF4, a TF more active in terminal interfollicular keratinocyte differentiation, top regulatory gene targets included soluble regulators of keratinocyte differentiation (DMKN and KRTDAP), as well as structural components of spinous and granular keratinocytes (KRT1,2 and IVL) (Figure 3K). Again, we found that these predicted KLF4 regulatory targets were enriched for genes that were downregulated (OR = 1.87, Fisher’s exact test p-value = 4.9 x 10^-7^), but not upregulated (OR = 0.95) in keratinocytes upon KLF4 knockdown ^59^. These 387 putative KLF4 target genes were enriched for GO-term processes including cornification and establishment of skin barrier. We inspected the gene targets of a number of TFs involved in HF keratinocyte function as well, and identified distinct groups of gene targets associated with ‘cholesterol storage’ for AR, which is expressed in isthmus HF keratinocytes, and WNT-protein binding for LHX2, which is expressed in inferior segment HF keratinocytes (Figure S3F,G).

#### Preservation of HFSCs and depletion of bulbar HF keratinocytes in alopecia areata

Alopecia areata results in disruption of normal hair follicle cycling by auto-reactive cytotoxic T-lymphocytes. While we observed increased overall abundance of T-lymphocytes in both scATAC and scRNA samples from patients with alopecia areata (Figure 1H,I, S1I,J), we did not observe any discrete populations of T-lymphocytes unique to patients with alopecia areata, nor did we identify an appreciable number of differential accessible chromatin peaks between AA and control T-lymphocytes, perhaps because the cellular phenotype of auto-reactive T-lymphocytes in AA is relatively subtle, or is driven by especially rare subpopulations of cells that we could not identify. In our sub-clustered keratinocyte dataset, we found that select HF-associated keratinocyte populations appeared to be depleted in samples from patients with AA (Figure S4A). We used Milo^60^ to test for differential abundance of keratinocytes between AA and control samples, and confirmed that AA samples had relatively fewer cells corresponding to the inferior segment of the hair follicle (Inf.Segment)(Figure 4A,B). Upon further sub-clustering of cells that we anticipate originate in the inferior segment of the hair follicle, we identified six populations of HF keratinocytes in the bulbar and suprabulbar region of the hair follicle (Figure 4C, S4B-D). We annotated these as quiescent hair follicle stem cells (HFSCs: KRT15, CD200, LHX2, NFATC1)^43, 61–63^, two populations of sheath cells (Sheath_1/2: SOX9, KRT5, KRT75)^49^, a population of matrix cells (Matrix: LEF1, KRT81, KRT82, HOXC13)^64, 65^, and a population of hair germ cells (CD34, LGR5, CDH3, WNT3)^66^. We used Milo to test for differential abundance of these populations between AA and control samples and found that AA samples demonstrated relative preservation of HFSC abundance, but a depletion of inferior segment sheath populations (Figure 4E). The observed preservation of HFSCs and depletion of inferior segment sheath populations are consistent with the known non-scarring, relapsing and remitting nature of AA. These findings also support the theory that sheath cell populations in the hair bulb are especially affected by the disrupted immune environment in AA ^35, 67, 68^.

**Figure 4.**
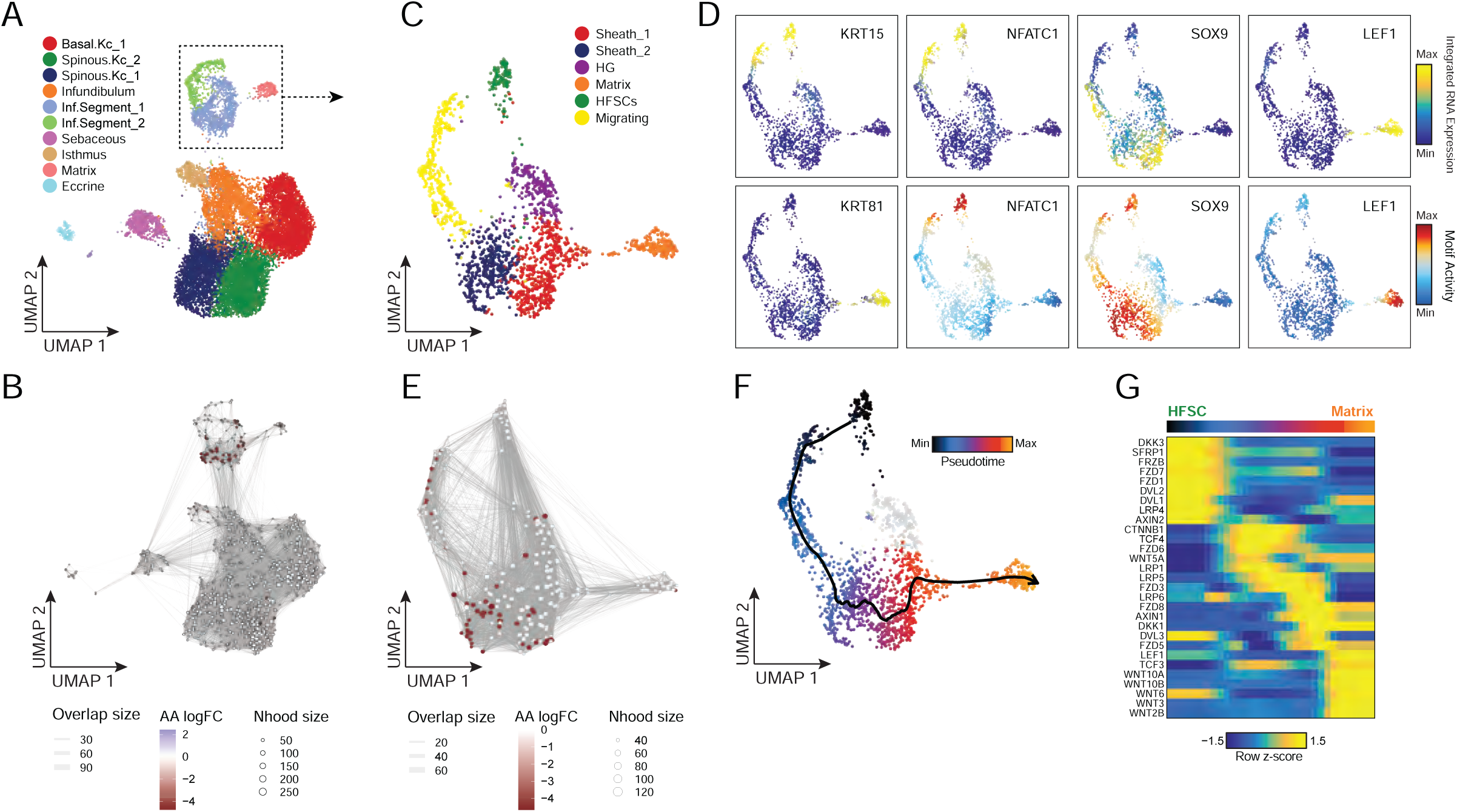
Regulatory dynamics of human hair follicle cycling (A) Sub-clustered keratinocytes in scATAC-seq space. The inferior segment of the hair follicle is highlighted. (B) Differential abundance of cycling HF keratinocytes between alopecia areata and control samples using Milo. Colored spots represent neighborhoods that are differentially abundant with a SpatialFDR < 0.1 (C) Sub-clustered HF keratinocytes from the inferior segment of the hair follicle. (D) Selected marker gene expression and TF motif activity deviation z-scores for sub-clustered inferior segment HF keratinocytes. (E) Same as in B, except for sub-clustered cycling HF keratinocytes. Hair sheath cells are differentially depleted relative to HFSCs. (F) Differentiation trajectory from HFSCs to Matrix cells. (G) Heatmap of variable expression of members of the WNT signaling pathway during HF cycling.

#### WNT-pathway dynamics through *in vivo* human HF keratinocyte differentiation

The WNT signaling pathway plays an essential role in both hair follicle development, post-natal hair follicle cycling, and in hair follicle regeneration after wounding^40, 52, 69–72^. However, the majority of studies of WNT pathway activity in hair follicle cycling have come from *in vitro* or mouse *in vivo* systems. To explore the dynamics of WNT signaling in human hair follicles, we constructed a semi-supervised pseudo-temporal trajectory from quiescent HFSCs to cycling HF matrix cells (Methods) (Figure 4F). We correlated ChromVAR TF motif activity with corresponding TF expression using cells along this pseudotime trajectory to identify putative drivers of this differentiation process (Figure S4E). Consistent with previous studies in mice, we found that NFATC1 was active in quiescent HFSCs, and as differentiation continued the WNT associated TFs TCF3 and TCF4 became expressed and active in sheath cells, and LEF1 became expressed and active in matrix cells ^61, 73, 74^. We performed GO term enrichment on the top 10% of variable expressed genes across this trajectory and observed significant enrichment of genes from the WNT-signaling pathway (Figure S4F). To explore the dynamics of WNT signaling over the course of HFSC differentiation, we plotted the expression of WNT signaling factors and downstream receptors across pseudotime (Figure 4G). We found HFSCs robustly expressed WNT receptors FZD1 and 7, but also several soluble WNT inhibitors (DKK3, SFRP1) and the soluble FZD receptor FRZB, suggesting that these cells may be primed to respond to paracrine WNT signaling, but maintain quiescence by actively blocking these signals. As HF keratinocytes progress through the differentiation trajectory, expression of WNT-pathway inhibitory signals decreases and the expression of beta-catenin (CTNNB1) and WNT activated TFs (TCF3 and TCF4) increases. Consistent with previous genomic studies in mice, actively dividing matrix cells in the hair bulb express multiple activating WNT effectors (WNT3, WNT5A, WNT10A/B) and the TF LEF1 becomes more highly expressed and active (Figure 4H, S4E)^40, 75^.

#### Cell-type specific open chromatin regions in scalp are enriched for skin and hair GWAS traits

Most complex skin and hair diseases are highly polygenic, and the vast majority of identified variants associated with disease are in non-coding regions of the genome^4, 6, 8, 76^. To determine if particular cell types in human scalp are more likely to be involved in the pathoetiology of skin and hair disease, we used cell-type specific open chromatin regions to perform linkage disequilibrium (LD) score regression using GWAS studies of skin, hair, and non-skin related traits (Methods, Figure 5A, S5A,B)^77, 78^. We observed strong enrichment of androgenetic alopecia (AGA) per-SNP heritability across fibroblast cluster open chromatin regions, with the strongest enrichment in dermal papilla (DP) peaks– the component of the hair follicle reported to have the highest androgen receptor activity ^79, 80^. We also found modest but significant SNP heritability enrichment for AGA in open chromatin regions of several HF keratinocyte clusters. Autoimmune skin diseases including psoriasis and atopic dermatitis had significant enrichment of SNP heritability in T-lymphocyte open chromatin regions, while tanning and hair pigment color were most enriched in melanocyte open chromatin regions. Traits not directly related to scalp cell types, such as Schizophrenia or BMI, did not demonstrate any cell type specific enrichment (Figure 5A).

**Figure 5.**
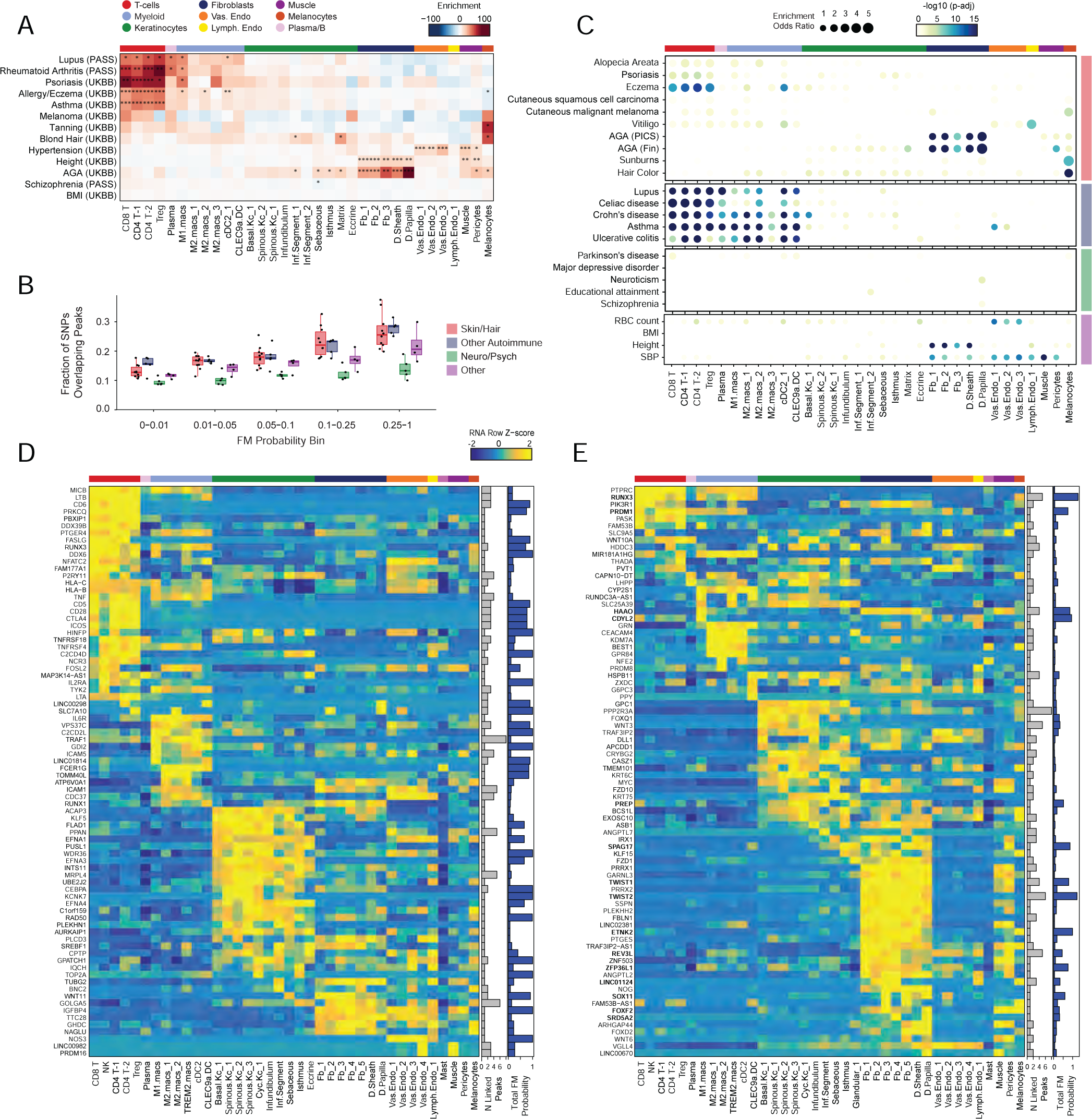
Identification of cell types and genes associated with hair, skin, and autoimmune diseases (A) LD score regression identifies enrichment of GWAS SNPs for various skin and non-skin related conditions in peak regions specific to sub-clustered cell types in human scalp. FDR-corrected P-values from LDSC enrichment tests are overlaid on the heatmap (*FDR < 0.05, **FDR < 0.005, ***FDR < 0.0005). (B) Fraction of fine-mapped SNPs overlapping scalp open chromatin regions binned by increasing fine-mapping posterior probability. Color of boxplot indicates the group of traits being plotted. Boxplots represent the median, 25th percentile and 75th percentile of the data, and whiskers represent the highest and lowest values within 1.5 times the interquartile range of the boxplot. (C) Fisher’s exact test enrichment for fine-mapped trait-related SNPs in peak regions specific to sub-clustered cell types in human scalp. The FDR-corrected -log_10_ p-value is indicated by the color of the dots, and the dot size indicates the enrichment odds ratio. Traits are grouped as in (B). (D) The top genes linked to peaks containing fine-mapped SNPs for eczema. The heatmap shows relative gene expression for each high-resolution scRNA cluster. The number of linked peaks per gene is indicated in the gray bar plot to the right, and the total sum of fine-mapped posterior probability for linked SNPs is indicated in the blue bar plot. (E) Same as in (D), but for androgenetic alopecia.

Because LD score regression requires full GWAS summary statistics, which were not available for several traits including AA, we also examined the enrichment of fine-mapped SNPs in cell type specific open chromatin regions^81–83^. We found that fine-mapped SNPs for skin, hair, and autoimmune disorders were more likely to overlap scalp cis-regulatory elements than fine-mapped SNPs for neurodegenerative and psychiatric disorders, and that this difference became greater with increasing fine-mapping posterior probability (Figure 5B, S5C). We again observed cell-type specific enrichment for several disease associated fine-mapped SNPs (Fisher’s exact test, FDR < 0.05) (Figure 5C). Alopecia Areata fine-mapped SNPs showed the most significant enrichment in CD4 T-cell (OR = 3.28, Fisher’s exact test adjusted p-value = 0.0013) and Treg (OR = 3.47, Fisher’s exact test adjusted p-value = 0.0058) open chromatin regions, but also were significantly enriched in several myeloid lineage clusters (e.g. M2.macs_2; OR = 2.80, Fisher’s exact test adjusted p-value = 0.0018). Interestingly, while height-associated fine-mapped SNPs were enriched in fibroblast clusters broadly, there was little to no enrichment of fine-mapped SNPs in the DP cluster, while AGA fine-mapped SNPs were most strongly enriched in DP open chromatin regions (OR = 5.58, Fisher’s exact test adjusted p-value = 1.3 x 10^-32^, Figure 5C).

#### Integrated transcriptomic and epigenetic single cell data enables linkage of fine-mapped SNPs with potential target genes

After nominating disease-relevant cell types in the scalp, we next sought to identify the genes with expression influenced by the presence of fine-mapped GWAS variants (GWAS target genes). For a given disease or trait, we aggregated the posterior probability of fine-mapped SNPs lying in linked peaks for each gene (Methods). We then plotted the expression of each of the top 80 genes by aggregate fine-mapped posterior probability across all high resolution scRNA clusters in the full scalp dataset to identify cell type-specific expression of genes linked to fine-mapped SNPs. For eczema, we identified a total of 137 genes linked to fine-mapped SNPs (fmGWAS-linked genes), the majority of which were expressed in T-cell or keratinocyte clusters (Figure 5D). These genes included important modulators of immune signaling such as TNF, CTLA4 and FASLG, and previously nominated GWAS gene targets including IL6R, PUS10 and IL2RA ^8, 84^. For androgenetic alopecia, we identified 130 fmGWAS-linked genes, the majority of which were accessible in keratinocyte or fibroblast clusters (Figure 5E). These fmGWAS-linked genes were enriched for TFs (OR = 3.18, Fisher’s exact test p-value = 4.1 x 10^-7^), such as TWIST1, TWIST2, RUNX3, FOXF2, FOXD2, and SOX11, and included several members of the WNT signaling pathway such as WNT3, WNT6, WNT10A, FZD1, and FZD10. GO-term enrichment of the top 80 AGA fmGWAS-linked genes also revealed enrichment of terms related to the WNT-signaling pathway (Figure S5D). For alopecia areata, we identified only 31 fmGWAS-linked genes in our dataset, likely due to the less highly powered GWAS available for this trait, but these included notable genes involved in T-cell functions such as IL21, ICOS and IRF4 (Figure S5E). IL21 in particular had multiple fine-mapped SNPs residing in linked enhancer elements, is expressed most specifically in CD4 T-cells, has been shown to play a role in supporting the persistence of cytotoxic CD8 T-cells in chronic viral infections^85, 86^ and has been implicated in the etiology of several autoimmune diseases ^87^. For hair color, we identified 158 fmGWAS-linked genes, the majority of which were expressed in keratinocyte subpopulations and melanocytes (Figure S5F).

#### Machine learning models prioritize potential functional noncoding SNPs for skin and hair phenotypes

In addition to nominating cell types and gene targets associated with skin and hair disease, we sought to obtain yet higher-resolution information by identifying SNPs that exert their effects by modulating TF binding and enhancer function. To prioritize fine-mapped SNPs that might directly alter TF binding, we implemented a gapped k-mer support vector machine (gkm-SVM) learning framework to score the allelic effect of a SNP on cell-type specific chromatin accessibility, a proxy for differential TF binding ^88–91^. We trained predictive models using the scATAC-seq data from 28 scalp clusters that had sufficient numbers of cells for training (Methods) (Figure 6A). These models demonstrated high and stable predictive performance on held-out test sequences using a 10-fold cross validation scheme (Figure S6A-D). We used GkmExplain to predict the per-base impact of specific variants in a given target sequence in each cluster by providing each model with sequences containing both the reference and alternative allele for a candidate SNP^92^. To create a set of prioritized SNPs associated with AGA, eczema, and hair color, we selected SNPs that (1) had a fine-mapping posterior probability ≥ 0.01, (2) overlapped scalp cis-regulatory elements, and (3) were predicted to disrupt chromatin accessibility in our model. We found that prioritized SNPs for eczema were enriched in multiple keratinocyte and T-cell clusters relative to random fine-mapped SNPs residing in scATAC-seq peaks, while prioritized SNPs for hair-color were enriched more specifically in hair follicle associated keratinocytes and melanocytes (Figure 6B, S6E). We did not observe notable cluster-specific enrichment of AGA prioritized SNPs, perhaps due to the specificity of this trait for dermal papilla cells and the lack of a dermal papilla-specific model given the low number of these cells in our dataset (Methods) (Figure S6F). We next filtered these prioritized SNPs to include only those that were linked to a target gene using our peak-to-gene linkage analysis, increasing the interpretability of potential causative variants. Using these criteria, we identified 47, 19, and 19 prioritized SNPs for AGA, eczema, and hair color respectively.

**Figure 6.**
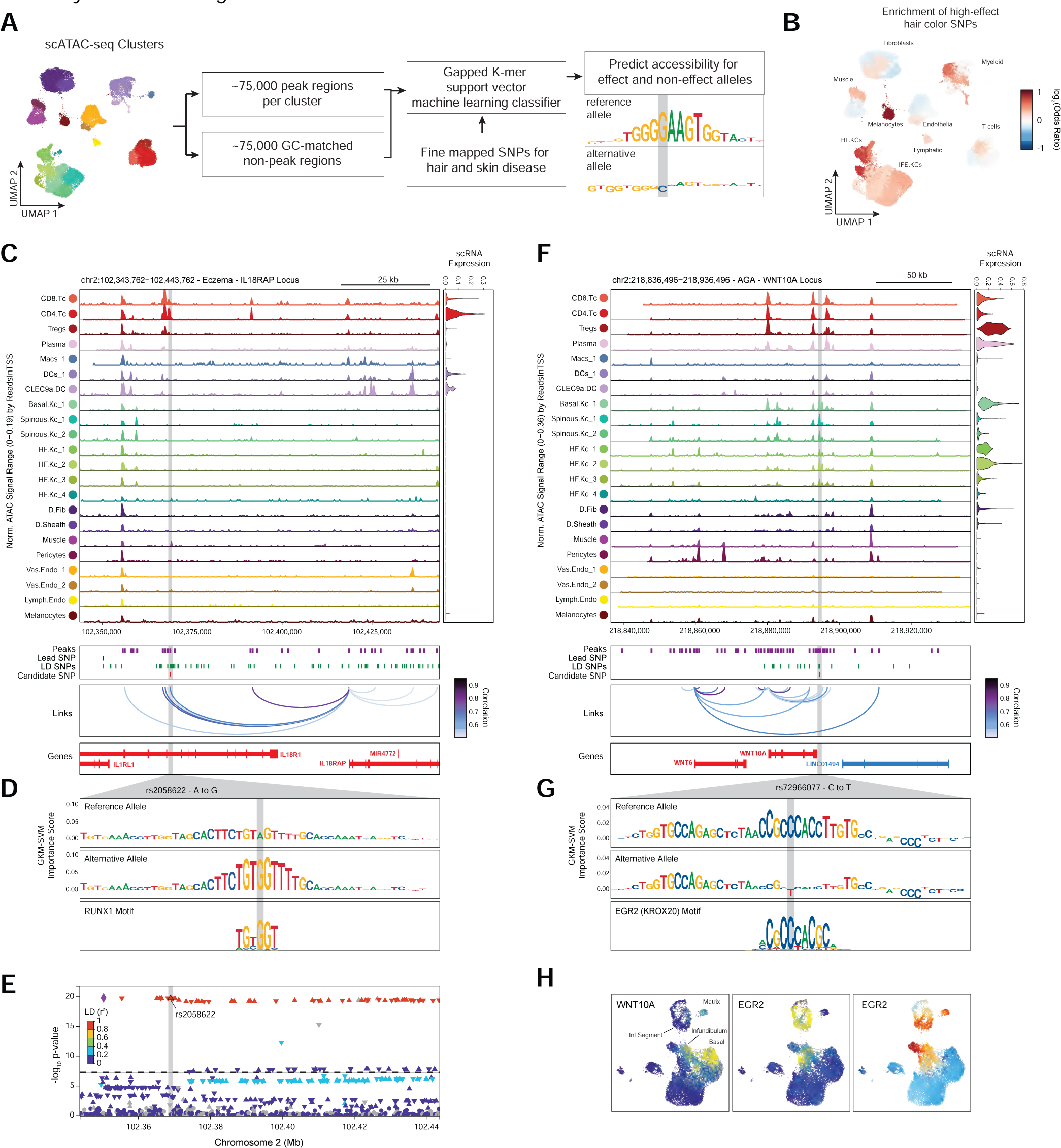
Candidate causal variants in skin and hair disease (A) Schematic of strategy for identification of potential causative variants. (B) Enrichment of high-effect fine-mapped SNPs from select skin and hair traits relative to random fine-mapped SNPs in cis-regulatory regions. (C) Normalized chromatin accessibility landscape for cell type-specific pseudo bulk tracks around the IL18RAP locus. Integrated IL18RAP expression levels are shown in the violin plot for each cell type to the right. The position of ATAC-seq peaks, the GWAS lead SNP, the fine-mapped SNP candidates in LD with the lead SNP, and the candidate functional SNP are shown below the ATAC-seq tracks. Significant peak-to-gene linkages are indicated by loops connecting the IL18RAP promoter to indicated peaks. (D) GkmExplain importance scores for the 50bp region surrounding rs2058622, an eczema associated SNP that disrupts a RUNX motif in a cis-regulatory element linked to IL18RAP expression. The effect and non-effect alleles for the gkm-SVM model correspond to the model trained on the CD4 helper T-cell cluster. (E) LocusZoom plot of the region shown in (C), highlighting the strong LD and high overall GWAS signal of this locus. (F) Same as in (C), but for the WNT10A locus. (G) GkmExplain importance scores for the 50bp region surrounding rs72966077, an androgenetic alopecia associated SNP that disrupts a ERG motif in a cis-regulatory element linked to WNT10A expression. The effect and non-effect alleles for the gkm-SVM model correspond to the model trained on the Infundibular keratinocytes cluster. (H) UMAP projection of the high-resolution keratinocyte subclustering showing expression of WNT10A, EGR2, and EGR2 ChromVAR motif activity.

One of the high-effect SNPs we identified for eczema was rs2058622, which is located in an intron of the IL18R1 gene (Figure 6C). This candidate SNP overlapped a cis-regulatory element that was preferentially accessible in the CD4 helper T cell cluster, and although this cis-regulatory element was within an IL18R1 intron, we found that this peak was linked to IL18RAP expression in our peak-to-gene linkage analysis. Our CD4 T cell base-resolution model suggested that the alternative allele of this SNP increases cell-type specific chromatin accessibility at this peak by creating a strong RUNX motif (Figure 6D). Furthermore, this SNP had been previously identified as a highly significant eQTL for IL18RAP expression in blood, with the G allele increasing expression (p-value 4.8 x 10^-54^, Normalized Effect Size (NES) 0.28). While this locus is one of the most strongly associated with eczema^8, 84^, it is also a region with significant linkage disequilibrium and contains multiple potential gene targets, making identification of causal SNPs for this locus challenging and highlighting the utility of our multi-tiered approach (Figure 6E). IL18RAP is an accessory protein that is required for potentiation of IL-18 signaling through the IL18 receptor complex ^93^, and IL-18 overexpression in the skin of mice induces a phenotype similar to atopic dermatitis ^94^, suggesting a mechanistic pathway for this causal variant.

One of the high-effect SNPs identified for androgenetic alopecia was rs72966077, which is located immediately downstream of the WNT10A gene body (Figure 6F). Interestingly, this SNP has also been implicated in acne vulgaris, another hair follicle and androgen associated disease^95^. This SNP overlapped a cis-regulatory element accessible in multiple keratinocyte clusters, although WNT10A expression was highest in basal keratinocytes and in infundibular hair follicle keratinocytes. Our infundibular keratinocyte model demonstrates that the alternative allele of this SNP disrupts an ERG family transcription factor motif (Figure 6G). Examination of the expression of all ERG family transcription factors across keratinocytes revealed that ERG2 is expressed in infundibular, isthmus, and inferior segment hair follicle keratinocytes, and these cell populations also have more ERG2 ChromVAR motif activity compared to other keratinocyte populations (Figure 6H). Patients with naturally occurring WNT10A mutations exhibit numerous skin appendage-related phenotypes, including a pattern of hair thinning that resembles androgenetic alopecia^96^. Furthermore, depletion of ERG2 (also known as Krox20) positive hair-follicle keratinocytes in mice results in arrest of hair growth^97^. These converging lines of evidence highlight the importance of the WNT-signaling pathway in the pathobiology of AGA and show that even though the strongest AGA GWAS signal enrichment is in dermal papilla cells, there may also be keratinocyte-intrinsic genetic factors that contribute to this complex trait.

## Discussion

In this study, we generated high-resolution, paired epigenomic and transcriptomic atlases of primary human scalp, a complex tissue harboring dynamic and precisely regulated hair follicles. By integrating these orthogonal datasets across multiple levels of cellular resolution, we identified principles of variable gene expression across diverse cell types, defined gene regulatory networks across interfollicular and hair follicle associated keratinocytes, and used a tiered approach to prioritize cell types, genes, and causal variants implicated in the pathobiology of skin and hair associated phenotypes.

By identifying peak-to-gene linkages at multiple levels of cellular resolution, we were able to capture both broad, cell-class regulatory differences, as well as regulatory programs delineating more subtle cell type differences. This approach yielded 50-100% more enhancer-gene links than similarly powered single-cell studies, even with a more conservative correlation threshold for calling linked peaks^17, 40, 41^. Using an orthogonal dataset of enhancer-gene predictions, we showed that these additional enhancer-gene linkages identified on sub-clustered datasets were confirmed with similar frequency to those identified on the full dataset (Figure S2I).

Several HRGs, linked to a disproportionately large number of cis-regulatory elements, are expressed in multiple distinct cell types, and the aggregate accessibility of computationally linked, cell-type specific enhancer modules predicts the resulting level of expression (Figure 2F-I). These findings support a broadly additive model of enhancer activity on gene expression, where the magnitude of expression is proportional to integrated effect of multiple individual, and generally interchangeable, cis-regulatory elements. We posit that this regulatory strategy makes expression of genes related to a cell’s core function more resistant to perturbation, but also allows for exquisitely tunable expression of a gene across multiple cellular contexts. This mode of enhancer activity is consistent with a recent high-throughput reporter-assay study of enhancer-promoter interactions that demonstrated that the variability in intrinsic enhancer activity was low compared to the target promoter’s intrinsic activity, and that enhancer elements are generally interchangeable in their ability to drive expression from a given promoter^48^.

Sub-clustering of our keratinocyte cellular compartment enabled identification of differentiation pseudotime trajectories for both interfollicular and hair-follicle associated keratinocytes. We used our integrated chromatin and transcriptomic data to nominate key transcription factors active in both steady state keratinocyte subpopulations and at different points along the IFE keratinocyte differentiation trajectory. To identify potential regulatory targets of these TFs, we used an analytical framework that takes into account the correlation between a TF’s global motif activity and a potential target gene, as well as the presence of linked enhancer elements that contain instances of that TFs binding motif. This approach allowed us to confirm key roles for TP63 in coordinating hemidesmosome assembly, and KLF4 in promoting upper layer keratinocyte differentiation progression and cornification. For most of the TFs investigated, high linkage scores for a gene target were associated with a clear positive correlation between global TF motif activity and target gene expression (Figure 3I,K, S3F,G), but for some TFs, such as FOSL1 and POU2F3 (Figure 3J, S3H), there were several gene targets with high linkage scores but negative correlation to global TF motif activity. This may imply a negative regulatory role for these TFs for select gene targets. Indeed, selective transcriptional repression has been described previously for both FOSL1^98^ and POU2F3^99, 100^. These findings implicate important target genes and regulatory networks for further investigation and treatment of hair-associated traits and disease.

Our analyses using LDSC and enrichment of fine-mapped SNPs in open chromatin regions provided meaningful insights into driver cell types for multiple hair and skin diseases. While we see significant GWAS signals for AGA in some HF keratinocyte subpopulations, by far the most significant enrichment of GWAS signal was in dermal papilla open chromatin regions (Figure 5). This genomic prioritization is consistent with functional studies showing that DP cells have the most robust AR expression among HF cell types^79^, and exhibit differential gene expression and signaling behavior when isolated from balding and non-balding individuals^101, 102^. Unsurprisingly, GWAS signals for autoimmune diseases were highly enriched in immune cell types, but these signals of enrichment were often quite cell-type specific– for example, diseases with prominent autoantibody pathologies such as lupus and rheumatoid arthritis showed enrichment in plasma cells, while cell-mediated autoimmune diseases, such as eczema, did not. Interestingly, even highly skin-specific autoimmune diseases such as eczema and psoriasis with clear keratinocyte phenotypes clinically, showed little enrichment for GWAS signals in keratinocyte subpopulations relative to T-lymphocyte cell types, suggesting that the genetic susceptibility to these diseases is primarily immunological, and less due to genetic variation intrinsic to keratinocytes.

Finally, using our multi-omic scalp datasets to train cell-type specific machine learning models of chromatin accessibility, we nominated potential functional SNPs for multiple skin and hair diseases. Our multi-pronged approach allowed us to trace the regulatory effect of single-base changes in the genome through their modulation of TF binding, to disruption of target gene expression in the relevant cell type in human scalp. We note, however, that for several strongly associated loci we were unable to identify a potential causal SNP target. This could be because the regulatory elements in which these SNP effects become apparent are only observed in the disease state, or the relevant pathologic cell type is not present in the tissue investigated. Indeed, DP cells were most enriched for SNP heritability for AGA (Figure 5A,B), but because of their rarity, we were unable to train a cell-type specific model, thus limiting our ability to more precisely predict regulatory effects in these cells. Moreover, several traits may also be the end result of a developmental process, and the relevant regulatory regions may be dormant in adult tissues. We anticipate that future studies will be able to surmount these challenges as the costs of single-cell sequencing continue to decrease, tissue collection methods improve, and models of gene regulation continue to be refined.

## Acknowledgments

We thank the Stanford FACS facility and Stanford Functional Genomics Facility for technical support. Figure schematics were created with BioRender.com. This work was supported in part by grants from the NIH 2R37-ARO54780 (AEO), RM1-HG007735 (WJG), UM1-HG009442 (WJG), UM1-HG009436 (WJG), R01-HG00990901 (WJG), and U19-AI057266 (WJG.). W.J.G. acknowledges support as a Chan-Zuckerberg Investigator (grant # 2017-174468 and 2018-182817). B.O.R. was supported in part by the Stanford MSTP training grant (T32GM007365). C.W. was supported in part by a Dermatology Foundation Physician Scientist Career Development Award.

## Author Contributions

Conceptualization, B.O.-R., C.W., A.E.O. and W.J.G.; Investigation-Single Cell Experiments, B.O.-R. and C.W.; Formal Analysis, B.O.-R.; Resources-Sample Collection, C.W., A.E.O., J.M.K., E.J.R. and S.Z.A..; Writing – Original Draft, B.O.-R. and W.J.G.; Writing – Review and Editing, All Authors; Supervision, A.E.O., M.M.D., W.J.G.; Funding Acquisition, A.E.O., M.M.D, and W.J.G.

## Methods

### Sample acquisition and patient consent

Primary human scalp samples were obtained in the form of 4mm punch biopsies or from excess discarded scalp tissue from patients undergoing dermatological surgeries (surgical “dogears”). Samples were collected from Stanford University or from Santa Clara Valley Medical Center with Stanford University Institutional Review Board approval, and all patients provided written informed consent. Upon collection, samples were stored in 1x PBS at 4℃ until dissociation and downstream processing. Samples were stored no longer than 5 h prior to dissociation. A 4mm punch was performed on surgical dogear samples prior to proceeding with sample dissociation.

### Single Cell Dissociation and Fluorescent Activated Cell Sorting (FACS)

Scalp punch biopsies were rinsed with ice cold 1x PBS and then lightly diced into 1-2mm pieces with a sterile razor blade. The diced sample was then dissociated using the Miltenyi Biotec Human While Skin Dissociation Kit (Product no. 130-101-540) according to the manufacturer’s directions. Briefly, samples were incubated in 0.5 mL dissociation solution containing the indicated volumes of Enzymes P, A, and D for 3 h at 37℃. Following incubation, 0.5 mL of ice cold RPMI 1640 w/ 10% FBS was added to each sample and then samples were mechanically dissociated using the gentleMACs dissociator with the ‘h_skin_01’ program. Following dissociation, samples were briefly centrifuged and then filtered through a 70µm cell strainer. The dissociation tube was washed with additional ice cold media and then samples were centrifuged for 10 minutes at 300 x g in a swinging bucket centrifuge. After aspirating the supernatant, samples were either resuspended in 0.1mL BamBanker freezing medium (Wako Chemicals, 302-14681) and cryopreserved at -80℃, or proceeded immediately to staining for FACS. We did not observe any systematic clustering differences between samples that had been sorted immediately after dissociation or those that had been cryopreserved, even without Harmony or other batch correction methods (Figure S1B,C). Cryopreserved samples included C_SD4, C_SD5, C_SD6, C_SD7, AA7, and AA8. All remaining samples were sorted immediately after dissociation without cryopreservation.

Cells were stained with anti-CD90 PE Cy7 (BD Pharmingen, order no. 561558) for 30 min at 4℃ in FACS staining buffer (PBS with 0.5% BSA), then washed with FACS buffer. Live cells were distinguished using LIVE/DEAD Fixable Aqua Dead Cell Stain Kit (ThermoFisher, order no. L34957) according to the manufacturer’s directions. For cryopreserved samples, cells were thawed at 37℃ for 3 min, resuspended in RPMI + 10% FBS and washed with FACS buffer prior to staining. Aqua negative live cells were sorted as fibroblast (CD90+) and non-fibroblast (CD90-) populations. Sorted cells were counted and the CD90+ population was reduced by half before recombining with the CD90-population for further processing by single cell ATAC-seq and/or single cell RNA-seq.

### Bulk ATAC-seq of subset of control samples

Bulk ATAC seq was performed on dissociated cells from four of the surgical dogear control samples (C_SD4, C_SD5, C_SD6 and C_SD7). These samples were thawed quickly in a 37℃ water bath and then 1 mL of prewarmed media (RPMI 1640 w/ 10% FBS) was added to each sample. Samples were centrifuged at 300 x g for 5 minutes at 4℃ and the supernatant was aspirated. Each sample was resuspended in ice cold 1x PBS with 0.5% BSA and then split into two aliquots of 200,000 cells. If fewer cells were present for a sample, the number of cells was split evenly. One aliquot from each sample was pooled and this pool was used for two lanes of 10x single cell ATAC-seq v2 (see below). The remaining aliquot of each sample was used for bulk ATAC-seq, which was performed similarly to as described previously^103^. Briefly, cell aliquots were centrifuged at 300 x g for 5 minutes at 4℃ and the supernatant was aspirated.

### Each pellet was resuspended in 100 µL of ice cold lysis buffer (10 mM Tris-HCl, pH 7.4, 10 mM

NaCl, 3 mM MgCl2, 1% BSA, 0.01% Digitonin, 0.1% Tween-20, and 0.1% NP40) and incubated on ice for 3 minutes. The lysis reaction was then diluted by the addition of 1 mL of ice cold RSB-washout buffer (10 mM Tris-HCl, pH 7.4, 10 mM NaCl, 3 mM MgCl2, 1% BSA, 0.1% Tween-20). Samples were centrifuged at 500 x g for 10 minutes at 4℃. The supernatant was aspirated and each nuclei pellet was resuspended in 50 µL of Transposition solution (10 mM Tris-HCl, pH 7.4, 5 mM MgCl2, 10% dimethyl formamide, 0.33x PBS, 0.01% digitonin, 0.1% Tween-20, 100 nM Illumina Tn5 Transposase (Illumina, 20034197)). Samples were incubated at 37℃ in a thermomixer rotating at 1000 RPM for 30 minutes. The remainder of the bulk ATAC-seq library generation was performed as described previously. Resulting bulk-ATAC libraries were pooled and sequenced on an Illumina NextSeq 550 using paired-end 36-bp reads.

### Genotyping and sample deconvolution using demuxlet

Bulk ATAC-seq FastQ files were processed and aligned to the hg38 reference genome as described previously^104^. Peaks regions identified from these bulk ATAC-seq samples were genotyped and used as input for Demuxlet to identify the patient sample for each cell for the pooled scATAC-seq samples as described previously^18,105^.

### Single Cell RNA library generation, sequencing, and alignment

Following sorting, cell suspensions were centrifuged at 300 x g for 5 minutes at 4℃ and resuspended in 1x PBS with 0.5% BSA. Samples were counted using a hemocytometer and the required volume of cells was aliquoted for generation of single-cell RNA sequencing libraries.

Single-cell RNA-seq libraries were prepared using the 10x Genomics Chromium Next GEM Single Cell 3′ RNA v3.1 protocol, targeting 8,000 cells per sample. Completed libraries were sequenced on an Illumina NextSeq 550 platform with 28/8/0/91 base pair cycles. Raw sequencing data was converted into fastq format using the command ‘cellranger mkfastq’ (10x Genomics, v3.1.0). Resulting fastq files were then aligned to the hg38 reference genome (cellranger-GRCh38-3.0.0) and quantified using the command ‘cellranger count’.

### Single Cell ATAC library generation, sequencing, and alignment

After the required number of sorted cells were aliquoted for generation of scRNA-seq libraries, the remaining sample volume was used for generation of single-cell ATAC sequencing libraries. The remaining cell volume was used to prepare nuclei according to the 10x ATAC nuclei isolation protocol for ‘low cell input nuclei isolation’ (CG000169, Rev B). Single cell ATAC-seq libraries were prepared using the 10x Genomics Chromium Next GEM Single Cell ATAC v1.1 protocol, targeting 6,000 cells per sample. Completed libraries were sequenced on an Illumina NextSeq 550 platform with 33/8/16/33 base pair cycles. Raw sequencing data was converted into fastq format using the command ‘cellranger-atac mkfastq’ (10x Genomics, v1.2.0). Resulting fastq files were aligned to the hg38 reference genome (cellranger-atac-GRCh38-1.2.0) and quantified using the command ‘cellranger-atac count’.

### scRNA-seq quality control, dimensionality reduction, and clustering

Following alignment and quantification, scRNA-seq count matrices were further processed using the ‘Seurat’ R package (v4.0.4) ^36^. Initial quality control was performed on each sample independently. First, cells were removed if they had less than 200 genes expressed, less than 1000 unique sequenced reads (UMIs), or greater than 20% of counts corresponding to mitochondrial genes. Doublets were identified and removed using the ‘DoubletFinder’ R package (v2.0.3) ^106^. Because we observed evidence of ambient RNA contamination in several samples, we used the ‘DecontX’ method in the ‘celda’ R package (v1.6.1) to estimate and remove contaminating ambient RNA from each cell ^107^. After each of these quality control steps were performed, samples were merged into a single Seurat object for clustering.

### Decontaminated count data was scaled to 10,000 and then log2-normalized

We adapted an iterative latent semantic indexing (LSI) approach to dimensionality reduction and clustering as described previously ^17^. First, we removed mitochondrial genes, sex chromosome genes, and genes associated with cell cycle (Seurat’s ‘cc.genes’) to minimize sample batch effects in variable feature selection. Next, we identified the top 4,000 variable genes across all cells and calculated the Term Frequency - Inverse Document Frequency (TF-IDF) for these variable genes. We performed Singular Value Decomposition (SVD) on the TF-IDF matrix and used the first 25 dimensions as input into Seurat’s SNN clustering with an initial resolution of 0.2. Counts from single cells in each of these resulting clusters were summed, transformed using a logCPM transformation ‘edgeR::cpm(mat, log=TRUE, prior.count=3),’ and then used to identify the top 4,000 variable genes for the next round of LSI. The TF-IDF transformation followed by SVD was performed again using the new set of 4,000 variable genes, and clustering was repeated with an increased resolution of 0.4. The previously described variable gene selection, TF-IDF transformation, and SVD was performed once more, and clustering was repeated with a final resolution of 0.8. The 25 LSI dimensions from the final round were used to generate 2-dimensional representations using the uniform manifold approximation and projection (UMAP) implementation from the Seurat and ‘uwot’ R packages (v1.0.10; n.neighbors=50, min.dist=0.5, metric=cosine).

This initial clustering procedure identified 29 clusters. After identifying marker genes for each cluster using Seurat’s ‘FindAllMarkers’ function and inspecting sample representation of each cluster, we identified a small number of clusters that appeared to be doublet clusters (Clusters 18, 26, 28, and 29). Each of these clusters were composed entirely or nearly entirely from a single sample, did not have unique marker genes when compared to other clusters, or expressed biologically incompatible combinations of marker genes. We removed all cells belonging to these clusters and repeated the previously described iterative LSI clustering procedure on the remaining cells, this time using clustering resolutions of 0.1, 0.3, and 0.6 for the three rounds. We regenerated the UMAP with the same parameters as used previously.

This final filtered and clustered dataset contained 21 clusters. Visualization of gene expression on UMAP representations was smoothed using the MAGIC diffusion algorithm ^108^. To minimize risk of ‘over smoothing’ expression patterns, the application of MAGIC was restricted to data visualization^109^.

#### scATAC-seq quality control, dimensionality reduction, and clustering

Following alignment, ATAC-seq fragment data was further processed using the ‘ArchR’ R package (v1.0.1) ^18^. For each cell, we computed the number of unique sequenced fragments and the transcription start site (TSS) enrichment, which serves as a signal to noise metric for ATAC-seq data^17^. We plotted all barcoded droplets on a scatter plot using these two metrics and observed that while some samples had a clear separation between true cells (high TSS and number of unique fragments), some samples had a more continuous distribution between true cells and droplets containing contaminating free DNA (lower number of unique fragments and lower TSS enrichment). To label droplets as likely true cells, we used an expectation maximization (EM) based approach. For each sample, we used the ‘mclust’ R package (v5.4.7) to fit up to 4, 2-dimensional gaussians to the log10 nFragments by TSS enrichment joint distribution (‘Mclust(df, G=2:4, modelNames=”VVV’). Cells classified as originating from the gaussian with the greatest mean TSS enrichment were labeled as true cells, while the remaining droplets were filtered from the project. Cells with a TSS of < 5 or nFragments < 1000 were all filtered from the project regardless of their EM classification label. This approach was functionally similar to setting a hard filter for TSS and nFragments for samples that had clearly defined true cell populations, but enabled exclusion of more contaminating droplets for samples that had a less clearly defined population of true cells (Figure S1A).

Following initial quality control, doublets were identified and filtered using the ArchR ‘addDoubletScores’ and ‘filterDoublets’ functions, with a filter ratio of 1. We then used ArchR’s implementation of iterative LSI dimensionality reduction using the ‘addIterativeLSI’ function with 50,000 variable features and 25 dimensions. We identified clusters using the ArchR function ‘addClusters’ with a resolution of 0.6, and then we generated a 2-dimensional representation of the data using the ‘addUMAP’ ArchR function with nNeighbors=50, minDist=0.4, and metric=cosine). This initial clustering procedure identified 22 clusters. We identified marker genes for each cluster using the ‘getMarkerFeatures’ function with the accessibility around each gene (the ‘Gene Activity Score’) as a proxy for gene expression^18^. We identified a small number of poor quality clusters (Clusters 7, 13, 15, and 18). These clusters did not have unique marker genes, had systematically lower TSS enrichment, or were enriched for high doublet scores. Cells belonging to these clusters were filtered from the project, and dimensionality reduction and clustering was repeated on the filtered project using 50,000 variable features and 50 dimensions for ‘addIterativeLSI’, and then a resolution of 0.7 for ‘addClusters’. We regenerated the UMAP using nNeighbors=60, minDist=0.6, and metric=cosine. This final filtered and clustered dataset contained 22 clusters. Visualization of gene activity scores on the UMAP was similarly smoothed using the MAGIC algorithm ^108^. Smoothed data was used only for visualization purposes as described previously.

#### Sub-clustering of major cell types

To improve identification of rare cell types, we sub-clustered several major cell groups from the full scRNA and scATAC-seq datasets. For scRNA-seq data, cluster labels were assigned based on known cell type markers (Figure 1E - ‘NamedClust’). Cluster labels for scATAC-seq data were assigned in a similar manner, using gene activity scores as a proxy for gene expression (Figure 1F - ‘NamedClust’). After labeling clusters in each modality, we sub-clustered major cell types in each dataset (Keratinocytes, Fibroblasts, Endothelial cells, T-cells, and Myeloid lineage cells) (Figure S2D-F). For scRNA-seq subclustering, we used the same iterative LSI dimensionality reduction procedure described above, except that we used two rounds of LSI instead of three, and we used the ‘Harmony’ R package (v0.1.0) to reduce sample batch effects since this effect was more pronounced on sub-clustered datasets relative to the full dataset ^110^. For both scRNA-seq and scATAC-seq subclustering, the number of variable features and the resolution of clustering was tuned to attempt to balance the number of identified clusters between modalities and to match the cellular heterogeneity of each sub-clustered group. For the sub-clustered scRNA-seq datasets, we used 2,000 variable genes and 15 SVD dimensions, and a clustering resolution of either 0.1 or 0.2 in the first round, followed by a clustering resolution of 0.3 in the final round. To generate UMAPs for each sub-clustered scRNA-seq dataset, we used n.neighbors=35 or 40, min.dist=0.4, and metric=cosine. For scATAC-seq subclustering, we again used ArchR’s implementation of iterative LSI dimensionality reduction, and we used Harmony to reduce sample batch effects. For each sub-clustered group, we used either 25,000 or 50,000 variable features, 25 dimensions, and between 0.2-0.4 resolution for clustering. To generate UMAPs for each sub-clustered scATAC-seq dataset, we used n.neighbors=35, min.dist=0.4, and metric=cosine. Following sub-clustering in each dataset, we assigned sub-cluster labels based on known marker genes (Figure S2D-F - ‘FineClust’).

#### Integration of scRNA- and scATAC-seq datasets

Starting with the full dataset, we matched each scATAC-seq cell with its closest corresponding scRNA-seq cell using a previously described multi-modal dataset integration technique based on canonical correlation analysis (CCA). Specifically, we used the ArchR function ‘addGeneIntegrationMatrix’, which uses Seurat’s ‘FindTransferAnchors’ function to integrate the datasets ^18,36^. We used nGenes=3,000 for integration of the full dataset. We computed the Jaccard index between scRNA- and scATAC-seq cluster labels of integrated meta-cells and observed high correspondence (Figure S2D,E). Furthermore, we identified the same major cell types in each dataset, with the exception of mast cells which were only observed in the scRNA-seq dataset (Figure S2E). We repeated this integration procedure for each of the previously described sub-clustered datasets (Keratinocytes, Fibroblasts, Endothelial cells, T-cells, and Myeloid lineage cells), using nGenes=2,000. For each sub-clustered dataset, we similarly observed high correspondence between scRNA and scATAC-seq derived cluster labels (Figure S2D).

#### Peak calling in scATAC-seq datasets

After sub-clustering of major cell types, we transferred the sub-cluster labels (‘FineClust’) back to the full scATAC-seq dataset for peak calling to maximize our ability to detect open chromatin regions specific to rare cell subtypes. Peak calling was carried out using the standard ArchR workflow. Pseudo-bulk group coverages were calculated for each cluster using the ArchR function ‘addGroupCoverages’, which were then used to call peaks using ‘addReproduciblePeakSet’. This function uses MACS2 (v2.1.1) to call fixed-width 500bp peaks on each cell type, merges the peak set from each cell type, and then iteratively removes overlapping peaks by dropping the lower scoring peak of overlapping pairs until no overlapping peaks remain ^18,104^. This procedure resulted in identification of 589,294 unique peaks for the entire dataset.

For each of the sub-clustered scATAC-seq datasets, only a relatively small subset of the full union peak set will be accessible in any of the cell types present. This results in excessive numbers of completely inaccessible regulatory regions being used for analyses on the sub-clustered datasets, which decreases statistical power. However, simply repeating peak-calling on each sub-clustered dataset would result in a peak set that does not represent a true subset of the full peak set due to the iterative overlapping peak removal procedure used to obtain the full union peak set. To obtain a peak set that is specific to a given sub-clustered dataset but is also a true subset of the full union peak set, we loaded the original peak calls for each cell type in the sub-clustered dataset and then kept only the subset of peaks from the union peak set that overlapped with peaks called on the sub-clustered cell types.

#### Linkage of gene regulatory elements to gene expression using integrated datasets

Cis-regulatory elements were linked to their potential gene targets (‘peak-to-gene links’) using a correlation-based approach ^104^. This procedure involves creating up to 500 partially overlapping pseudo-bulks of 100 k-nearest neighbors integrated single cells (‘low overlapping cell aggregates’). The peak counts of each pseudo-bulk are summed, as are the gene expression counts of the corresponding integrated scRNA-seq transcript profiles. Candidate peak-gene pairs are then identified by first associating peaks within a genomic distance of 250kb to the TSS of each gene, and then computing the Pearson correlation coefficient of the log2 normalized accessibility and gene expression counts. This procedure was carried out using the ‘addPeak2GeneLinks’ function in ArchR^18^. High confidence peak-to-gene links were obtained by retaining links that had a Pearson correlation coefficient of >0.5.

Because this correlation procedure is dependent on the dimensionality reduction of the particular dataset being used, and because the dimensionality reduction in turn is dependent on variable gene selection across the full dataset, we found that that using the entire scalp dataset for this analysis robustly identified peak-to-gene links that corresponded to regulatory interactions defining major cell types (e.g. keratinocytes vs T-cells), but was less efficient at recovering regulatory interactions between more closely related cell subtypes (e.g. specific hair follicle keratinocyte subsets) (Figure 2B). To increase our sensitivity for detecting peak-to-gene linkages distinguishing more fine-grained cell subtypes, we repeated the previously described peak-to-gene linking procedure on each sub clustered major cell type, using only the subset of peaks relevant to a specific sub-clustered dataset as described above. To create a consensus peak-to-gene link set, we combined all identified peak-to-gene links from the full dataset and each sub-clustered dataset, sorted peak-to-gene links by their Pearson correlation coefficients, and removed duplicate peak-to-gene links, resulting in a consensus peak-to-gene link set of 146,088.

#### Validation of inferred peak-to-gene linkages using conservation and ABC model predictions

Following identification of peak-to-gene linkages on the full scalp dataset and on each of the sub-clustered datasets (Keratinocytes, Fibroblasts, Endothelial, T-lymphocytes, and Myeloid), peak-to-gene links were validated using two strategies. First, we used the “gscores” function from the “GenomicScores” R package to compute the mean phastCons 100-way vertebrate evolutionary conservation scores for peaks linked in the full dataset and in each of the sub-clustered datasets, as well as for peaks that were not linked in any analysis. For each group of peak-to-gene linkages (i.e. the full dataset linkages and each of the sub-clustered datasets) we used a Wilcoxon rank-sum test to compare linked and unlinked peaks.

Second, we compared our peak-to-gene linkages to predicted enhancer-gene interactions from a recently published activity-by-contact (ABC) dataset generated from 131 human tissues and cell types ^38^. We downloaded the full dataset of all 131 tissues and cell types and converted enhancer coordinates from hg19 to hg38 using liftover. For validation of our peak-to-gene link inferences, we required both that the linked peak had to overlap an enhancer region in the ABC model dataset, and that the corresponding linked gene had to match. We used all possible peak-to-gene linkages (i.e. all peak-gene pairs separated by < 250 kb) as background to test for enrichment of ABC model predicted enhancer-gene links in our inferred peak-to-gene links (Figure S2I, top bar). To account for the skewed length distribution for inferred peak-to-gene links compared to all possible peak-to-gene links, we also compared the enrichment of ABC model predicted enhancer-gene links in inferred peak-to-gene links to a distance-matched background set of peak-to-gene links (Figure S2I, second bar). To do this, we first computed the distance between gene promoter and linked peak for all inferred peak-to-gene links. We divided these distances into 20 contiguous equal size bins and assigned background peak-to-gene links to each of these bins. We sampled 146,088 peaks from the background peak-to-gene link set while matching the distance distribution of the inferred peak-to-gene links, and then calculated the number of background peak-to-gene links that overlapped ABC enhancer-gene pair predictions. We repeated this sampling procedure 100 times, and used the mean number of overlapping background peak-to-gene links to calculate the enrichment of ABC enhancer-gene pair predictions in our inferred peak-to-gene linkages using a hypergeometric enrichment test. We calculated the enrichment of ABC model predicted enhancer-gene pairs in inferred peak-to-gene linkages for linkages identified on the full, non-sub-clustered dataset (“full scalp”), and for each of the sub-clustered datasets (Figure S2I, bottom 6 bars).

#### Identification and analysis of highly regulated genes (HRGs)

After creating our consensus peak-to-gene link set, we ranked all expressed genes by their number of peak-to-gene links. We found that a subset of genes had significantly more peak-to-gene linkages than others. We set a cutoff near the inflection point of 20 linked peaks per gene to identify a subset of ‘highly regulated genes’ (HRGs, 1,739 genes). We compared these HRGs to a dataset of previously identified super-enhancer associated genes from a variety of tissues and cell lines^42^. We also compared these HRGs to the human homologs of previously identified mouse hair-follicle associated super-enhancer genes ^43^. We calculated the enrichment of super-enhancer associated genes from various tissues in our set of 1,739 scalp highly regulated genes using a hypergeometric enrichment test (Figure 2D). In Figure 2E, we list two of the top HRGs for each k-means cluster to the right of the peak-to-gene heatmap.

#### Analysis of modular enhancer usage in HRGs

To visualize the heterogeneity of enhancer usage between cell types expressing the same gene, we generated 246 pseudo bulks of KNN cells with k = 250. To plot peak by pseudo bulk heatmaps, we normalized pseudo bulk accessibility by summing the peak counts for each pseudo bulk, depth normalizing and log2 transformed counts data, and then quantile normalization using the ‘normalize.quantiles’ function from the ‘preprocessCore’ R package. For each individual HRG, we then calculated the Z-score for the normalized accessibility of each linked peak across all pseudo bulk samples. Peaks were ordered using hierarchical clustering. For scatter plots comparing pseudo bulk linked peak accessibility to linked gene expression, we calculated the mean normalized integrated gene expression for each pseudo bulk sample and applied a log2 transformation. To calculate the total linked chromatin accessibility, we summed the depth normalized counts of linked peaks for a given gene and then applied a log2 transformation. Pseudo bulk labels in both the heatmaps and scatter plots were determined by selecting the most frequent cluster label from the 250 cells comprising each pseudo bulk.

#### ChromVAR motif analysis

We used chromVAR (v1.12.0) to measure the enrichment of transcription factor (TF) motifs in accessible chromatin across single cells ^111^. Specifically, we first used the ArchR function ‘addMotifAnnotations’ to identify all cisbp motif matches in the peak set, used ‘addBgdPeaks’ to identify a set of GC- and accessibility-matched background peaks, and then used the ‘addDeviationsMatrix’ function to calculate motif deviation Z-scores for each cisbp motif.

#### Trajectory analysis for interfollicular keratinocytes and hair follicle keratinocytes

To analyze epigenetic and gene regulatory dynamics over the course of differentiation of interfollicular keratinocytes, we used the R package ‘slingshot’ (v1.8.0) ^112^. To apply slingshot to our integrated scATAC-seq data for interfollicular keratinocytes, we used the ArchR function ‘addSlingShotTrajectories’ with ‘embedding=UMAP’, restricting available clusters to interfollicular keratinocyte clusters (Basal.Kc_1, Spinous.Kc_1, and Spinous.Kc_2), and designating the basal keratinocyte cluster as the origin of differentiation. To identify TF regulator candidates for this differentiation trajectory, we used two complementary approaches. First, using all keratinocyte clusters, we calculated the correlation between a given TF’s chromVAR motif deviation Z-scores and that same TF’s integrated gene expression across low-overlapping cell aggregates. Correlating these measures can help distinguish which specific TF in a larger TF family is responsible for the motif activity observed in a given cell type. These TF correlations were plotted against the maximum difference in chromVAR motif Z-scores between clusters, highlighting TFs exhibiting more dynamic regulatory activity across cell types (Figure S3E). To identify TFs that are more specific to the interfollicular keratinocyte differentiation trajectory, we selected integrated gene expression values and chromVAR deviation scores along the previously determined slingshot differentiation trajectory using the ArchR function ‘getTrajectory’ with groupEvery=1.5. We then correlated these trajectories using the ArchR function ‘correlateTrajectories’ with the default parameters.

To analyze the differentiation trajectory of the inferior segment of the hair follicle, we further sub-clustered these cells as described above. We used the keratinocyte scATAC-seq clusters Inf.Segment_1, Inf.Segment_2, and Matrix and scRNA-seq clusters Inf.Segment. For the sub-clustered scRNA-seq datasets, we used 1,500 variable genes and 20 SVD dimensions, and a clustering resolution of either 0.2 in the first round, followed by a clustering resolution of 0.4 in the final round. To generate UMAPs for the sub-clustered scRNA-seq dataset, we used n.neighbors=20, min.dist=0.1, and metric=cosine. For scATAC-seq subclustering, we again used ArchR’s implementation of iterative LSI dimensionality reduction. We used 25,000 variable features, 30 dimensions, and a 0.4 resolution for clustering. To generate UMAPs for the sub-clustered scATAC-seq data, we used n.neighbors=20, min.dist=0.1, and metric=cosine. We re-integrated these sub-clustered datasets and re-identified peak-to-gene linkages as described above. Note that this hair follicle inferior segment sub-clustering was used only for analysis of the hair follicle differentiation trajectory (Figure 4), and that the peak-to-gene links identified on this dataset were not used for any other analyses. Identification of TF regulators for the HF differentiation trajectory was performed using slingshot as described above, providing the HFSC, Migratory, Shaft_1, Shaft_2 and Matrix clusters as being involved in the trajectory and designating the HFSC cluster as the origin.

#### Identification of potential regulatory target genes of TF regulators

After identifying likely TF regulators as described above, we used the following strategy to identify potential regulatory gene targets of a given TF. First, we identified ∼500 low overlapping pseudo bulk samples of KNN with k=100 and obtained the mean normalized integrated gene expression and the mean chromVAR motif deviation score for each pseudo bulk sample. For our subset of candidate TF regulators, we used these pseudo bulk samples to calculate the Pearson correlation coefficient between the candidate TF regulator’s chromVAR motif activity and the integrated gene expression of all expressed genes. Next, we calculated a ‘Linkage Score’ for each gene and TF pair. This score is calculated by identifying all peak-to-gene links for that gene for which the linked peak contains an instance of the candidate TF motif, and then summing the product of the squared peak-to-gene linkage correlation with the the motif score:

[umath1]

Where LSg is the linkage score of gene g, n is the number of linked peaks for gene g, R is the peak-to-gene Pearson correlation coefficient for peak k, and MSk is the motif score for the motif occurring in peak k. The linkage score is thus higher for genes that have multiple linked peaks containing the TF motif, more strongly correlated linked peaks containing the TF motif, and/or linked peaks that contain highly confident instances of the motif. Finally, we also calculated the hypergeometric enrichment p-value for the TF motif in all linked peaks for a given gene. We defined potential gene regulatory targets of a TF regulator as those that have an absolute TF motif to gene expression correlation of >0.25 and a linkage score greater than the 80th percentile across all genes. We performed Gene Ontology (GO) enrichment analyses on the putative direct regulatory gene targets using the TopGO (v2.42.0) R package ^113^, using all expressed genes as background.

We validated inferred TF regulatory targets using previously published datasets of RNA-seq performed on keratinocytes with TF mutations or TF knockdown. For TP63, we downloaded the differentially expressed genes (Table S1D from Qu et al. 2018) identified between control human keratinocytes and keratinocytes containing a mutant, binding incompetent form of TP63^56^. We calculated the enrichment of downregulated genes from TP63 mutant keratinocytes in our predicted TP63 regulatory targets using a one-sided Fisher’s exact test. We also compared the enrichment of upregulated genes from these TP63 mutant keratinocytes in our predicted TP63 regulatory targets. For KLF4, we downloaded the raw, unnormalized counts matrix from a recent study that performed shRNA knockdown of KLF4 in human adult keratinocytes (GSE111786_counts_raw.csv.gz)^59^. We filtered genes that had fewer than 10 counts across all samples, and then performed normalization and differential testing using the DESeq2 R package (v1.30.1)^114^. We again calculated the enrichment of downregulated and upregulated genes from KLF4 knockdown keratinocytes in our predicted KLF4 regulatory targets.

#### Differential abundance testing using Milo

We used the ‘miloR’ R package (v1.1.0) to perform k-nearest neighbor graph-based differential abundance testing between alopecia areata (AA) and unaffected control samples (C_PB and C_SD) ^60^. While miloR was originally designed to be applied to scRNA-seq data, the algorithm depends only on having a cell-cell similarity structure to the dataset, and thus can be applied to scATAC-seq data. We applied miloR to our integrated scATAC-seq data by creating a ‘SingleCellExperiment’ R object from the counts matrix of our keratinocyte ArchR project and then used the ArchR LSI dimensionality reduction as the reduced.dim input for miloR in the ‘buildGraph’ function. For comparing differential abundance across all keratinocytes, we used only samples that had at least 50 cells in the sub-clustered dataset, and used k=30 for the ‘buildGraph’ function, and prop=0.1 for the ‘makeNhoods’ function. For comparing differential abundance across only the lower, cycling portion of hair follicle keratinocytes (Figure 4E), we used only samples that had at least 10 cells in the sub-clustered dataset, and used k=30 for the ‘buildGraph’ function and prop=0.3 for the ‘makeNhoods’ function. We plotted differentially abundant cell neighborhoods that had a SpatialFDR of < 0.1 using the ‘plotNhoodGraphDA’ function.

#### LD Score Regression using scATAC-seq data

We used linkage disequilibrium score regression (LDSR, v1.0.1) to estimate the heritability of multiple skin, hair and other traits in each high-resolution clustered cell type in our dataset ^78^. Briefly, this method determines if a functional category is enriched for heritability of a given trait by determining if SNPs with high LD to that category tend to have higher ꭓ^2^ statistics than SNPs with low LD to that category, conditioned on a baseline set of annotations. Cluster-specific peak regions were used as input functional categories for LDSR. To obtain these cluster-specific peaks, we first filtered clusters that had fewer than 40 cells total, as these clusters generally had too few cells to identify sufficient numbers of confident cell type specific peaks. For remaining clusters, we identified which peaks from the union peak set had been originally identified in a given cluster by overlapping the union peakset with the MACS2 peak calls from that specific cluster. For each cluster, we then retained only peaks that had been identified in no more than 25% of all clusters (9 out of a possible 36 clusters). This strategy enabled us to both filter out common ‘housekeeping peaks’ that are accessible in the majority of cell types, while retaining peaks that are unique to at most a few clusters. We formatted these cluster specific peaks using the ‘make_annot.py’ script, and LD scores were computed for each annotation using the ‘ldsc.py’ script with default parameters. We downloaded formatted summary statistics for partitioning from https://alkesgroup.broadinstitute.org/LDSCORE/all_sumstats/. We followed the recommended guidelines for cell-type-specific partitioned heritability analysis using the 1000G EUR phase 3 population reference and the hg38 baseline model (v2.2). We used the ‘ldsc.py’ script to calculate partitioned heritability for each trait in the cluster specific peak sets. We used Benjamini-Hochberg FDR correction to adjust heritability enrichment p-values.

#### Analysis of fine-mapped GWAS variants

We obtained fine-mapped SNPs from multiple sources. First, we downloaded a compendium of fine-mapped SNPs for 94 UKBB traits (www.finucanelab.org/data), and used the ‘Balding_Type4’, ‘BMI’ and ‘SBP’ traits for downstream analyses ^82^. Second, we downloaded pre-computed PICS fine-mapped SNPs for a variety of traits in the GWAS catalog (https://pics2.ucsf.edu/Downloads/PICS2-GWAScat-2021-06-11.txt.gz)^81,83^. We calculated enrichment of fine-mapped SNPs with a fine-mapping posterior probability of ≥0.01 from selected traits in the previously described cluster specific peak sets using one-sided Fisher’s exact test with a background SNP set containing all fine-mapped SNPs (also with a fine-mapping posterior probability of ≥0.01) across all traits. Enrichment p-values were adjusted using Benjamini-Hochberg FDR correction.

To identify genes associated with fine-mapped SNPs, for selected traits we identified fine-mapped SNPs that had a fine mapping posterior probability of ≥ 0.01 and overlapped a scATAC-seq peak region. Next, for each gene, we identified all fine-mapped SNPs that fell within a peak that was linked to the expression of that gene, and we summed the fine-mapping posterior probability for these linked SNPs. Genes linked to a peak containing a fine-mapped SNP with a high posterior probability, or genes linked to multiple linked peaks containing fine-mapped SNPs with appreciable fine-mapping posterior probability, are assumed to more likely represent genes whose expression is associated with the trait of interest. We plotted the row-scaled gene expression for the top 80 genes (by total associated fine-mapping probability) in each of our high-resolution scRNA-seq clusters in a heatmap, and plotted the number of linked peaks and the cumulative fine-mapping posterior probability to the right of each gene.

#### gkm-SVM machine learning classifier training and testing

We adapted a previously published strategy for trained gkm-SVM models using scATAC-seq data ^88^. For each scATAC-seq cluster, we trained a gapped k-mer machine learning classifier to predict whether a given genomic sequence is likely to be accessible or inaccessible in that cell type. We trained models for all clusters that had at least 200 cells to reduce training biases from spurious binding events from noisier, sparser data. We used training sequences of 1001 bp, expanding peaks on each side to reach this length and removing peaks that contained any N bases. As positive training data, we first identified the top 7,500 (by FDR) marker peaks for each cluster using the ArchR function ‘getMarkers’ with an FDR cutoff of 0.1 and a Log2FC cutoff of 0.5. These peaks represent the set of peaks that are most specific to a given cluster. We next identified which peaks from the union peak set had been originally identified in a given cluster by overlapping the union peakset with the MACS2 peak calls from that specific cluster. We sorted these by decreasing MACS2 score, and selected the top N peaks to combine with the previously identified marker peaks such that each cluster had ≤ 75,000 total unique peaks for training. For clusters that had fewer than 75,000 peaks, we used all peaks originally identified from that cluster as training peaks. Clusters that had fewer than 56,250 total peaks were not used for model training. To obtain negative training data, we generated 4,000,000 random genomic regions of 1001 bp (after masking assembly gaps (AGAPS) and intra-contig ambiguities (AMB)) and calculated the GC content of each of these regions. For each cluster, we then generated 20 equal-size bins of GC-content percentile from the N (≤ 75,000) positive training sequences. We labeled the random sequences according to these cluster-specific GC-content bins and sampled a total of N random regions while matching the binned distribution of GC-content from the original data.

After identifying the ≤ 75,000 positive and ≤ 75,000 negative training sequences for each cluster, we used a 10-fold cross-validation strategy to test model performance. We split training and testing sets by chromosome. The test sets for the 10 folds are as follows: Fold 1 consisted of chr 1; fold 2 consisted on chr 2 and 19; fold 3 consisted on chr 3 and 20; fold 4 consisted of chr 6, 13, and 22; fold 5 consisted of chr 5 and 16; fold 6 consisted of chr 4, 15 and 21; fold 7 consisted on chr 7, 14 and 18; fold 8 consisted of chr 11, 17 and X, fold 9 consisted of chr 9 and 12, fold 10 consisted of chr 8 and 10. For each fold, we used sequences from all non-testing chromosomes for training. For each of the 10 folds for each of the 29 clusters, we used the sequences from the positive and GC-matched negative training data as input training gkm-SVM models. Specifically, we used the LS-GKM package with the following options for the ‘gkmtrain’ function^91,115^. We used the wgkmrbf kernel (t = 5), a word length of 11 (l = 11), 7 informative columns (k = 7), up to 3 mismatches to consider (d = 3), an initial value of 50 for the exponential decay function (M = 50), a half-life parameter of 50 (H = 50), and a precision parameter of 0.001 (e = 0.001). We assessed the performance of trained models from cross-validation folds by calculating AUROC and AUPRC using the ‘PRROC’ R package (v1.3.1) with negative testing data downsampled to match the number of positive testing data sequences. To examine model specificity, we used the fold 0 from each cluster to predict the fold 0 testing data of every other cluster and again calculated the AUROC and AUPRC. Following assessment of model performance, we trained a full model for each cluster using all of the available training data.

#### Estimation of candidate SNP effect sizes using gkm-SVM models

We used our full gkmSVM models from each cell type to predict the change in accessibility for fine-mapped SNPs from GWAS for androgenetic alopecia (‘Balding_Type4’ from www.finucanelab.org/data and ‘Male-pattern baldness’ from PICS fine mapped), eczema (PICS fine mapped), hair color (PICS fine mapped), and 2500 randomly selected fine mapped SNPs overlapping ATAC peaks. We first selected only fine-mapped SNPs that overlapped a peak region, and then further filtered SNPs from all traits to those that had a fine-mapping posterior probability of ≥0.01. This resulted in 1,631 androgenetic alopecia SNPs, 612 hair color SNPs, 365 eczema SNPs, and 2500 random SNPs. For each SNP that met the above criteria, we obtained the 250bp surrounding the SNP and created synthetic alternative allele sequences by replacing the reference allele at the center of the sequence with the SNP alternative allele. We then used the previously trained full models for each cluster to calculate cell type-specific GkmExplain importance scores for each base of both the reference and alternative allele sequences for each of the fine-mapped SNPs^92^. GkmExplain estimates the per-base contribution for an input sequence to the corresponding output prediction of a gkmSVM model. We used GkmExplain for mutation impact scoring instead of DeltaSVM, ISM, or SHAP because it tended to yield more directly interpretable importance scores at the motif level, and has been previously shown to correlate extremely well with these other metrics ^88^. For each SNP and each cluster model, we summed the GkmExplain importance scores for the central 50bp of both the reference and alternative allele sequences, then subtracted the alternative allele score from the reference allele score to get the ‘delta score’ for the sequence immediately surrounding the SNP.

#### Statistical significance and prioritization of candidate SNPs

We used multiple metrics derived from GkmExplain importance scores to obtain a statistical significance for each tested fine-mapped SNP. First, for each SNP, we generated 3 di-nucleotide shuffled sequences using the ‘fasta-dinucleotide-shuffle.py’ script from the MEME suite ^116^. For each of these shuffled sequences, we generated a ‘reference’ and ‘alternative’ allele sequence corresponding to the original SNP by replacing the central position of the shuffled sequence with the SNP’s reference or alternative allele base. We calculated GkmExplain importance scores for each of these shuffled sequences across all clusters as described above, and calculated the cluster-specific ‘delta score’ using the central 50 bp of each null reference and alternative allele pair. The delta scores from these shuffled sequences served as a null distribution for each cluster. In agreement with a similar previous analysis, we found that the t-distribution was a good fit for these GkmExplain delta score null distributions ^88^. We used the ‘fitdistrplus’ R package (v1.1.6) to fit a t-distribution to each cluster’s delta score null distribution and used these distributions to calculate p-values for each fine-mapped SNP in each cell cluster. To further prioritize SNPs that are more likely to be affecting predicted accessibility by disruption of a transcription factor binding site, we calculated another previously described metrics of SNP effect size, the ‘prominence score’^88^. To calculate this score, we first identified the ‘active’ allele for each SNP by the sign of the delta score, with a positive delta score indicating that the reference allele is more likely to be accessible than the alternative allele. We then identified the subsequence surrounding the active allele SNP where each position’s GkmExplain importance score exceeded the 97.5th percentile of the di-nucleotide shuffled background. Subsequence boundaries were determined by the position where two consecutive bases had importance scores falling below the threshold. If a SNP sequence did not contain a subsequence of at least 7 bases, we used the central 7 bases surrounding the active allele as the subsequence. We used these subsequences to calculate the prominence score for each SNP. To calculate the prominence score, we took the sum of non-negative GkmExplain importance scores from the active allele subsequence and then divided by the sum of the non-negative importance scores for the entire 250bp sequence. This score can be thought of as a measure of the signal to noise ratio of the active allele for each SNP. We fit an exponential distribution to the prominence null distribution for each cluster and again used these distributions to calculate prominence p-values for each fine-mapped SNP in each cell cluster. To prioritize SNPs that have significant effects on predicted chromatin accessibility (large absolute delta score), likely through disruption of a transcription factor binding site (large prominence scores), we used the estimated p-values of these two scores determined from the shuffled sequence null distributions. We selected “high-effect” fine-mapped SNPs that had both a delta score p-value < 0.05, and had a prominence score p-value < 0.05. To increase interpretability and further filter for likely causal SNPs, we further filtered “high-effect” SNPs by requiring that they fall in a peak linked to expression of a gene. Using these criteria, we identified 47, 19, and 19 prioritized SNPs for AGA, eczema, and hair color respectively.

#### Data and software availability

Sequencing data has been deposited in the Gene Expression Omnibus (GEO) with the accession code GSE212450. Custom code for data processing, peak-to-gene analyses, and GWAS analyses is available on (https://github.com/GreenleafLab/scScalpChromatin).

---

**Figure S1.**
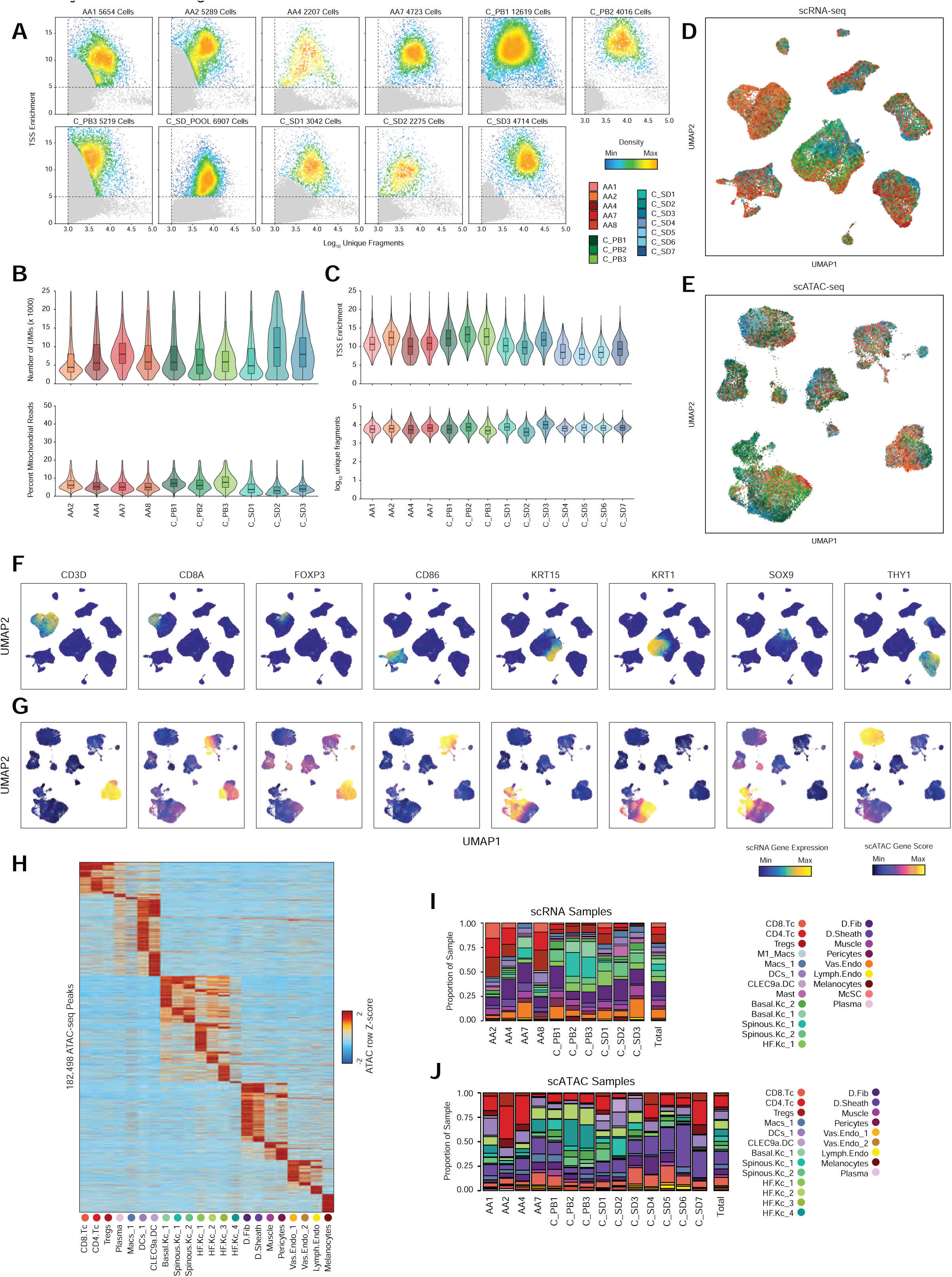
Quality control of single cell RNA and ATAC datasets. Related to Figure 1 (A) Scatter plots of the number of unique fragments by the transcription start site (TSS) enrichment for each of the scATAC-seq samples. Gray dots indicate cells that did not pass quality control filters (Methods). Colorbar indicates the density of points. (B) Violin plots of the number of unique reads (UMIs, top) and the percent of reads from mitochondrial genes (bottom) for each of the scRNA-seq samples.The inset box plot represent the median, 25th percentile and 75th percentile of the data, and whiskers represent the highest and lowest values within 1.5 times the interquartile range of the boxplot. (C) Violin plots of the TSS enrichment (top) and number of unique fragments (bottom) for each of the scATAC-seq samples. Box plot as in (B). (D) UMAP projection of full scRNA-seq dataset, colored by patient sample. (E) UMAP projection of full scATAC-seq dataset, colored by patient sample. (F) UMAP projections of full scRNA-seq dataset colored by relative expression levels of representative cell compartment marker genes. (G) UMAP projections of full scATAC-seq dataset colored by relative gene activity scores of the same marker genes shown in (F). (H) Marker peaks (Wilcoxon FDR ≤ 0.1 and Log2 fold change ≥ 0.5) for each scATAC cluster. (I) The fraction of each scRNA-seq cluster comprising each sample. The total proportions for each cluster are shown in the rightmost column. (J) The fraction of each scATAC-seq cluster comprising each sample. The total proportions for each cluster are shown in the rightmost column.

**Figure S2.**
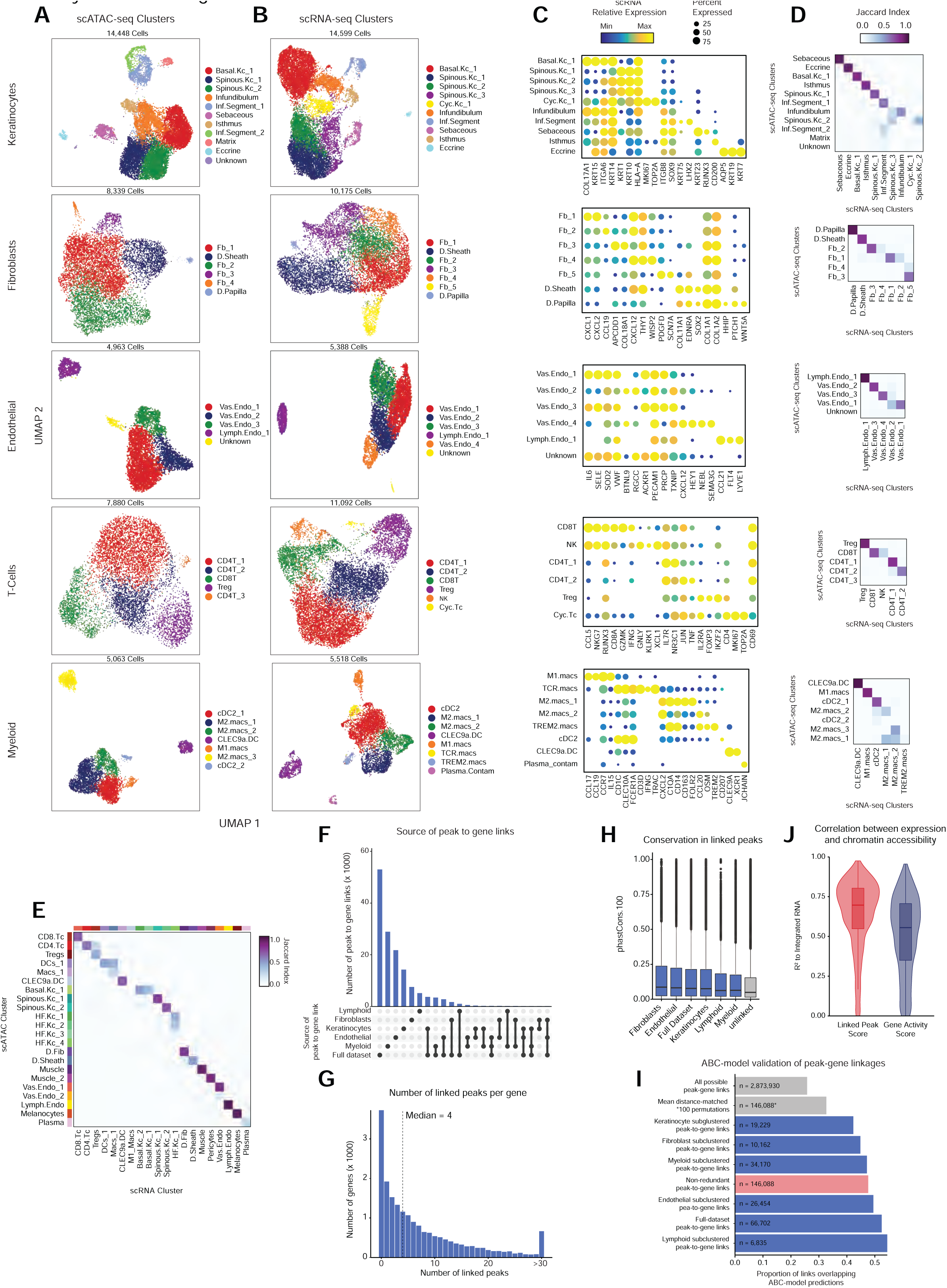
Sub-clustering of major cell groups, integration of scRNA and scATAC datasets, and identification of peak-to-gene linkages. Related to Figures 1 and 2. (A) UMAP representations of sub-clustered major cell groups using scATAC data. Cell compartments are labeled on the right, and cells are colored according to their high-resolution cluster labels. (B) UMAP representations of sub-clustered major cell groups using scRNA data. Cell compartments are labeled on the right, and cells are colored according to their high-resolution cluster labels. (C) scRNA gene expression for selected marker genes for each high-resolution scRNA-seq cluster from each sub-clustered cell group. The color indicates the relative expression across all high-resolution clusters and the size of the dot indicates the percentage of cells in that cluster that express the gene. (D) Correspondence between scRNA and scATAC-seq cluster labels for high-resolution clusters in each of the sub-clustered datasets. (E) Correspondence between scRNA and scATAC-seq cluster labels in the full scalp dataset. (F) Upset plot indicating the number of peak-to-gene linkages identified in the full dataset and in each of the sub-clustered datasets. (G) The distribution of the number of linked peaks per gene (median = 4). (H) The PhastCons 100-way vertebrate conservation scores for peaks with a linked gene in each dataset compared to unlinked peaks. Wilcoxon rank-sum test comparing each dataset to unlinked peaks, p < 2.2 x 10^-16^. Boxplots represent the median, 25th percentile and 75th percentile of the data, and whiskers represent the highest and lowest values within 1.5 times the interquartile range of the boxplot. (I) Bar plot showing the proportion of peak-to-gene linkages where both peak and gene were validated by a multi-tissue dataset of activity-by-contact (ABC) model enhancer-gene predictions. Categories compared included the space of all possible peak-to-gene links, the mean of 100 permutations drawn from all possible peak-to-gene links where for each permutation 146,088 peaks were selected to match the anchor distance distribution of true peak-to-gene links, and the set of true peak-to-gene links identified on each sub-clustered dataset. Hypergeometric enrichment tests comparing each subgroup of true peak-to-gene links to the mean distance-matched background set, p < 2.2 x 10^-16^. (J) Comparison of the linked peak score (sum of accessibility at linked peaks) compared to the gene activity score for predicting gene expression for the 1739 HRGs. Plotted is the Pearson R^2^ from 246 pseudo-bulked samples per gene. Boxplots represent the median, 25th percentile and 75th percentile of the data, and whiskers represent the highest and lowest values within 1.5 times the interquartile range of the boxplot.

**Figure S3.**
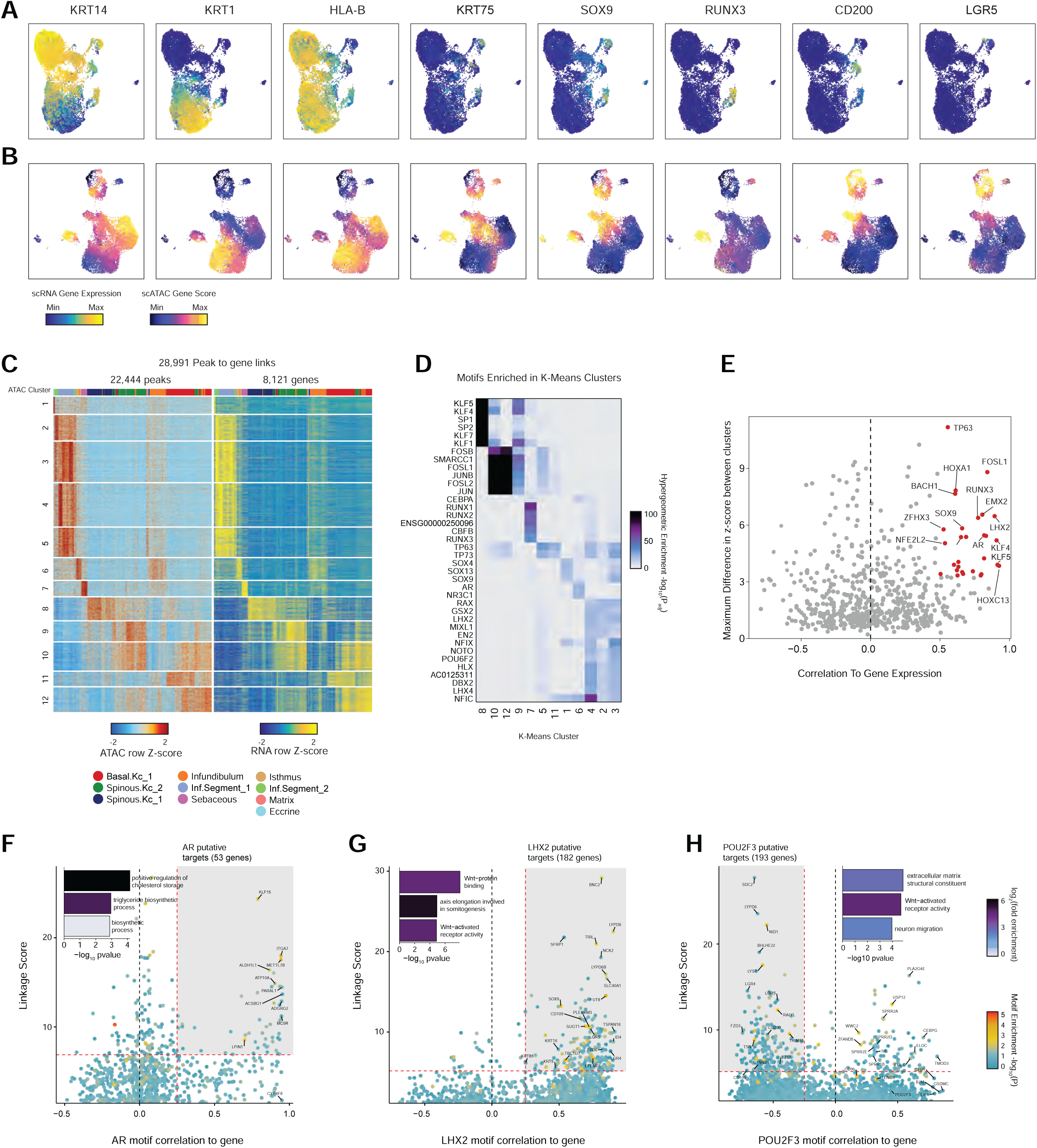
Marker genes and cell type specific TF regulator activity for sub-clustered interfollicular and hair-follicle associated keratinocytes. Related to Figure 3. (A) UMAP projections of sub-clustered keratinocyte scRNA-seq dataset colored by expression levels of representative marker genes. (B) UMAP projections of sub-clustered keratinocyte scATAC-seq dataset colored by gene activity scores of the same marker genes shown in (A). (C) Heatmap showing the chromatin accessibility (left) and gene expression (right) for 28,991 keratinocyte-specific peak-to-gene linkages. Peak-to-gene linkages were clustered using k-means clustering (k = 12). Rows indicate peak accessibility and gene expression on the left and right heatmaps respectively. Each column is a pseudo-bulk sample, with the colorbar on top of each heatmap indicating the cluster identity of each pseudo-bulk sample. (D) Hypergeometric enrichment p-values of TF motifs in peaks from each of the k-means clusters from (C). (E) Plot of TF motif activity correlation to corresponding TF gene expression across sub-clustered dataset against the maximum difference in chromVAR deviation z-score between clusters. TF’s with a maximum chromVAR difference in the top quartile and a pearson correlation greater than 0.5 are colored in red. (F) Prioritization of gene targets for LHX2. The x-axis shows the Pearson correlation between the TF motif activity and integrated gene expression for all expressed genes across all keratinocytes. The y-axis shows the TF Linkage Score (for all linked peaks, sum of motif score scaled by linkage correlation). Color of points indicates the hypergeometric enrichment of the TF motif in all linked peaks for each gene. Top gene targets are indicated in the shaded area (motif correlation to gene expression >0.25, linkage score >80th percentile). GO term enrichments for the top gene targets are shown in the inset bar plot. (G) Same as in (F), but for androgen receptor (AR). (H) Same as in (F), but for POU2F3.

**Figure S4.**
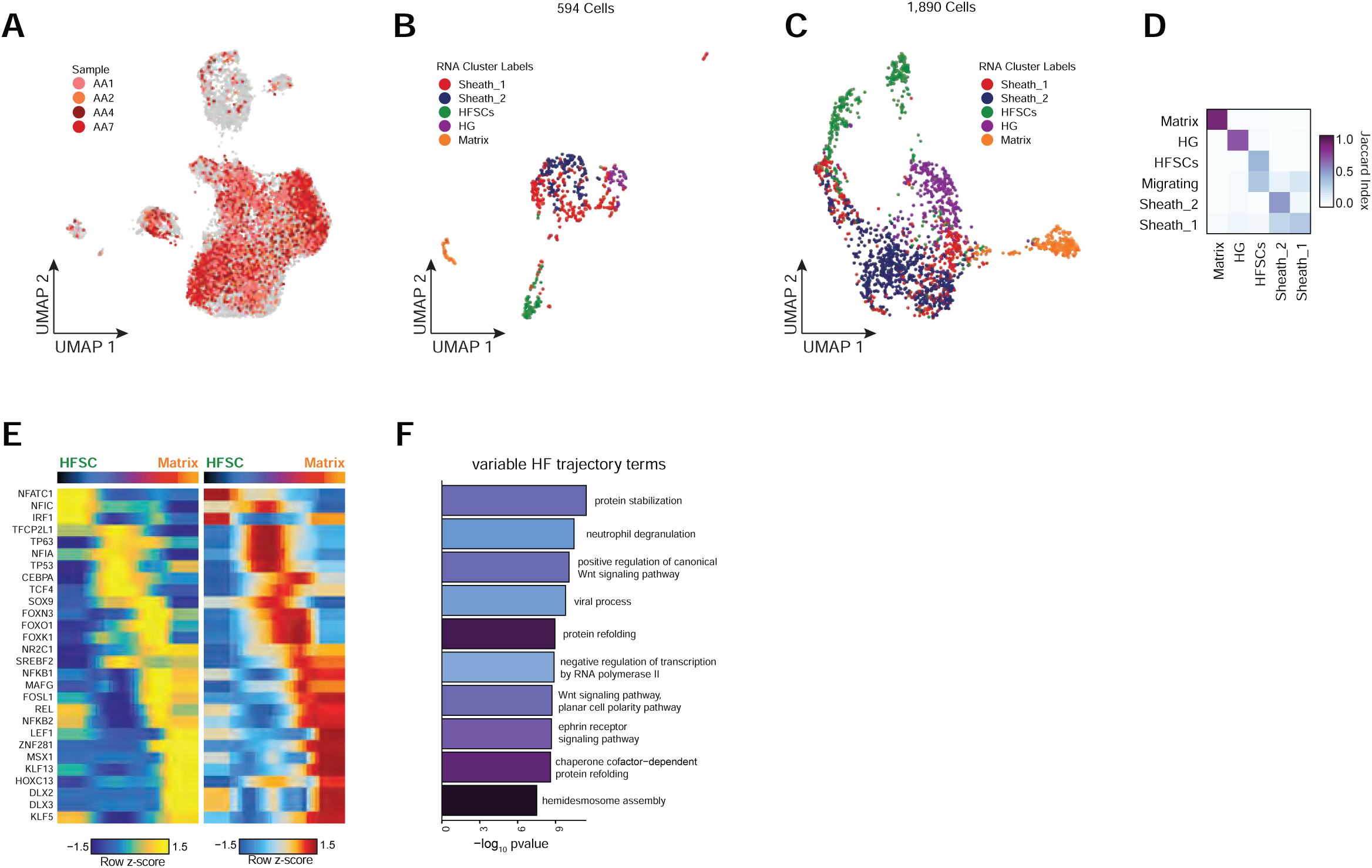
Supplemental analyses of sub-clustered inferior segment hair follicle keratinocytes. Related to Figure 4. (A) UMAP projection of sub-clustered keratinocytes showing cells originating from alopecia areata. Cells originating from control samples are colored gray and sorted to the back of the plot. (B) UMAP projection of sub-clustered scRNA inferior segment hair follicle keratinocytes. (C) UMAP projection of sub-clustered scATAC inferior segment hair follicle keratinocytes colored by matched nearest scRNA cluster. (D) Correspondence between scRNA and scATAC-seq cluster labels for integrated inferior segment hair follicle keratinocytes. (E) Paired heatmaps of positive TF regulators whose TF motif activity (left) and matched gene expression (right) are positively correlated across the hair follicle keratinocyte differentiation pseudotime trajectory. (F) GO term enrichments of the most variable 10% of genes across the hair follicle keratinocyte differentiation pseudotime trajectory.

**Figure S5.**
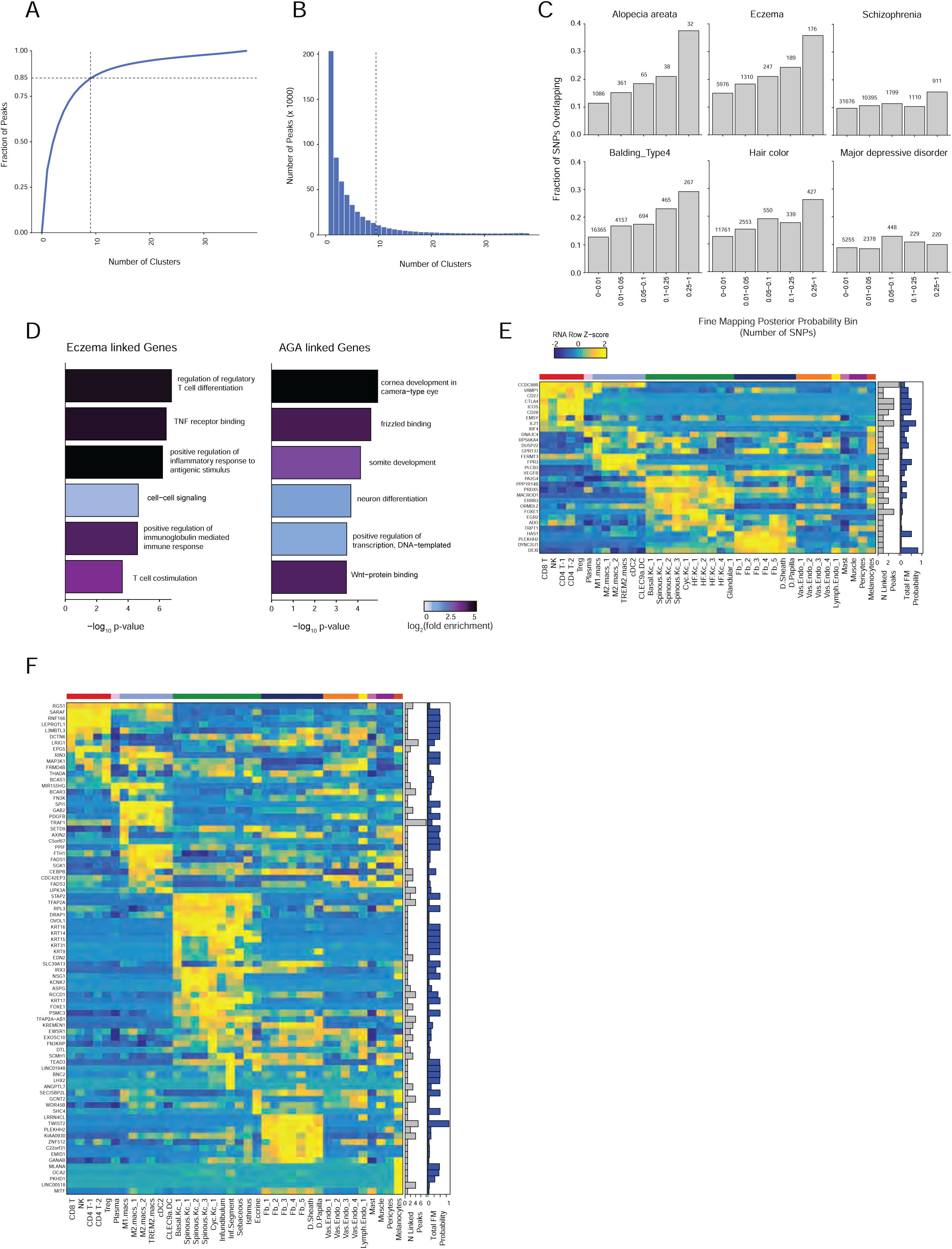
Supplemental analyses of fine-mapped SNPs overlapping scalp cis-regulatory elements. Related to Figure 5. (A) Cluster-specificity of peaks used for LD score regression and for fisher enrichment tests in Figure 5. More than 50% of peaks are specific to <1/8 of high-resolution scATAC clusters, and 85% of peaks are specific to ≤ 1/4 of clusters. (B) Distribution of the number of clusters in which each peak is accessible. Peaks accessible in ≤ 1/4 of clusters (9 high-resolution clusters) were used for cluster-specific enrichment analyses. (C) Fraction of fine-mapped SNPs for selected traits overlapping scalp cis-regulatory elements binned by fine-mapping posterior probability. (D) GO term enrichment for the top genes linked to fine-mapped SNPs by summed fine-mapping posterior probability in associated peak-to-gene linkages. (E) The top genes linked to peaks containing fine-mapped SNPs for alopecia areata. The heatmap shows relative gene expression for each high-resolution scRNA cluster. The number of linked peaks per gene is indicated in the gray bar plot to the right, and the total sum of fine-mapped posterior probability for linked SNPs is indicated in the blue bar plot. (F) Same as in (E), but for hair color.

**Figure S6.**
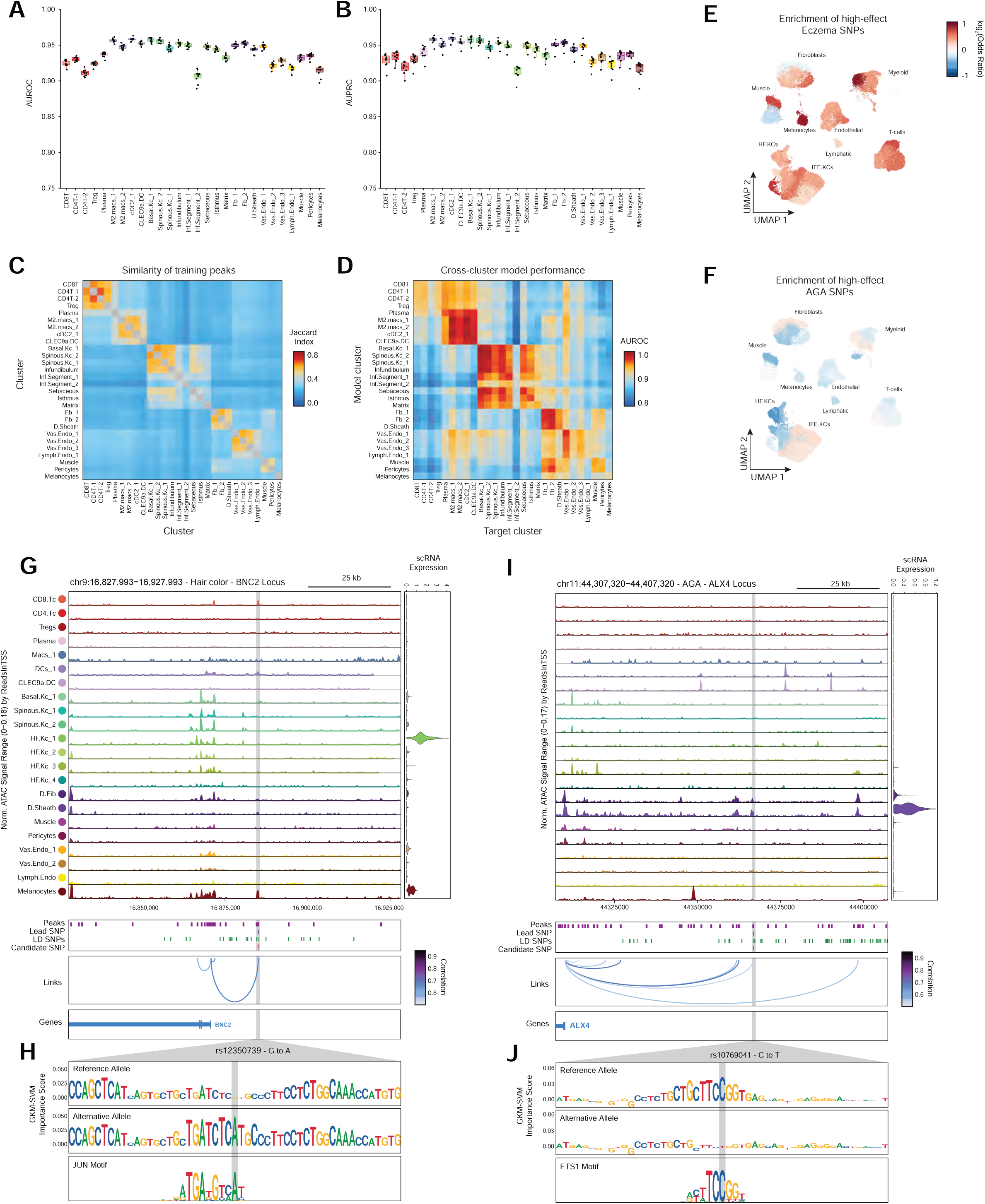
Assessment of gkmSVM model performance and additional high-effect candidate fine-mapped SNPs. Related to Figure 6. (A) The area under the receiver operator (AUROC), or (B) precision recall (AUPRC) curves for the gkm-SVM machine learning classifiers for each of the cluster models. Boxplots represent the median, 25th percentile and 75th percentile of the data, and whiskers represent the highest and lowest values within 1.5 times the interquartile range of the boxplot. (C) The overlap of training data (peak sequences) between models. (D) The performance of each cluster model on predicting test sequences from a non-target cluster. (E) Enrichment of high-effect fine-mapped SNPs from eczema relative to random fine-mapped SNPs in cis-regulatory regions. (F) Same as in (E), but for androgenetic alopecia. (G) Normalized chromatin accessibility landscape for cell type-specific pseudo bulk tracks around the BNC2 locus. Integrated BNC2 expression levels are shown in the violin plot for each cell type to the right. The position of ATAC-seq peaks, the GWAS lead SNP, the fine-mapped SNP candidates in LD with the lead SNP, and the candidate functional SNP are shown below the ATAC-seq tracks. Significant peak-to-gene linkages are indicated by loops connecting the BNC2 promoter to indicated peaks. (H) GkmExplain importance scores for the 50bp region surrounding rs12350739, a hair color associated SNP that creates a JUN motif in a cis-regulatory element linked to BNC2 expression. (I) Same as in (G), but for the ALX4 locus. (J) GkmExplain importance scores for the 50bp region surrounding rs10769041, an androgenetic alopecia associated SNP that disrupts an ETS motif in a cis-regulatory element linked to ALX4 expression.

## List of supplementary tables

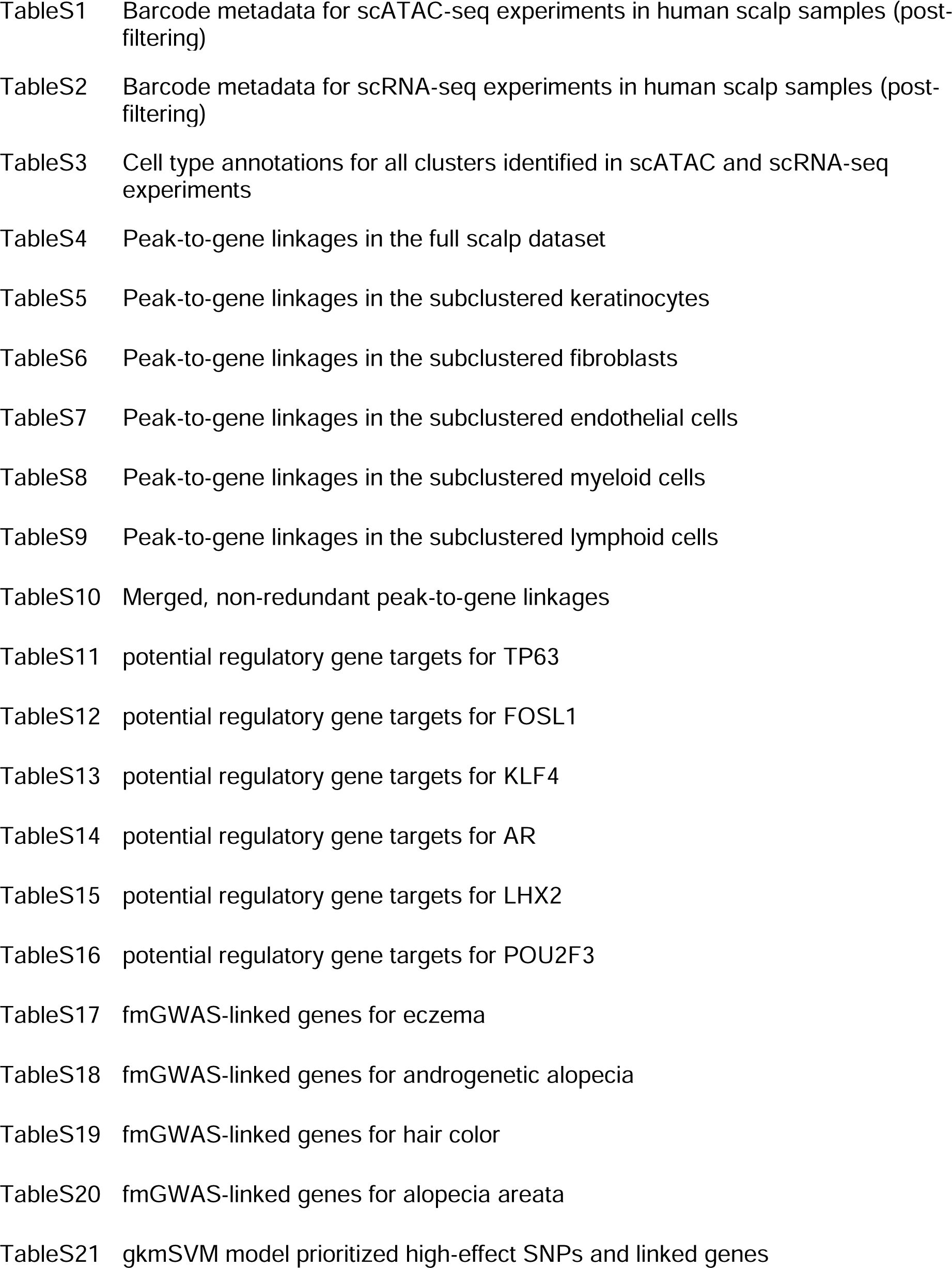

|  |  |
| --- | --- |
| TableS1 | Barcode metadata for scATAC-seq experiments in human scalp samples (post-filtering) |
| TableS2 | Barcode metadata for scRNA-seq experiments in human scalp samples (post-filtering) |
| TableS3 | Cell type annotations for all clusters identified in scATAC and scRNA-seq experiments |
| TableS4 | Peak-to-gene linkages in the full scalp dataset |
| TableS5 | Peak-to-gene linkages in the subclustered keratinocytes |
| TableS6 | Peak-to-gene linkages in the subclustered fibroblasts |
| TableS7 | Peak-to-gene linkages in the subclustered endothelial cells |
| TableS8 | Peak-to-gene linkages in the subclustered myeloid cells |
| TableS9 | Peak-to-gene linkages in the subclustered lymphoid cells |
| TableS10 | Merged, non-redundant peak-to-gene linkages |
| TableS11 | potential regulatory gene targets for TP63 |
| TableS12 | potential regulatory gene targets for FOSL1 |
| TableS13 | potential regulatory gene targets for KLF4 |
| TableS14 | potential regulatory gene targets for AR |
| TableS15 | potential regulatory gene targets for LHX2 |
| TableS16 | potential regulatory gene targets for POU2F3 |
| TableS17 | fmGWAS-linked genes for eczema |
| TableS18 | fmGWAS-linked genes for androgenetic alopecia |
| TableS19 | fmGWAS-linked genes for hair color |
| TableS20 | fmGWAS-linked genes for alopecia areata |
| TableS21 | gkmSVM model prioritized high-effect SNPs and linked genes |

